# Semi-synthetic CoA-α-Synuclein Constructs Trap N-terminal Acetyltransferase NatB for Binding Mechanism Studies

**DOI:** 10.1101/2023.04.03.535351

**Authors:** Buyan Pan, Sarah Gardner, Kollin Schultz, Ryann M. Perez, Sunbin Deng, Marie Shimogawa, Kohei Sato, Elizabeth Rhoades, Ronen Marmorstein, E. James Petersson

**Author notes:** **Corresponding Author** E. James Petersson - Department of Chemistry; University of Pennsylvania, Philadelphia, Pennsylvania 19104, United States.

## Abstract

N-terminal acetylation is a chemical modification carried out by N-terminal acetyltransferases (NATs). A major member of this enzyme family, NatB, acts on much of the human proteome, including α-synuclein (αS), a synaptic protein that mediates vesicle trafficking. NatB acetylation of αS modulates its lipid vesicle binding properties and amyloid fibril formation, which underlies its role in the pathogenesis of Parkinson’s disease. Although the molecular details of the interaction between human NatB (hNatB) and the N-terminus of αS have been resolved, whether the remainder of the protein plays a role in interacting with the enzyme is unknown. Here we execute the first synthesis, by native chemical ligation, of a bisubstrate inhibitor of NatB consisting of coenzyme A and full-length human αS, additionally incorporating two fluorescent probes for studies of conformational dynamics. We use cryo-electron microscopy (cryo-EM) to characterize the structural features of the hNatB/inhibitor complex and show that, beyond the first few residues, αS remains disordered when in complex with hNatB. We further probe changes in the αS conformation by single molecule Förster resonance energy transfer (smFRET) to reveal that the C-terminus expands when bound to hNatB. Computational models based on the cryo-EM and smFRET data help to explain the conformational changes and their implications for hNatB substrate recognition and specific inhibition of the interaction with αS. Beyond the study of αS and NatB, these experiments illustrate valuable strategies for the study of challenging structural biology targets through a combination of protein semi-synthesis, cryo-EM, smFRET, and computational modeling.

## INTRODUCTION

N-terminal acetylation is a ubiquitous modification made to proteins, with most of the eukaryotic proteome and ∼80% of the human proteome bearing this feature.^1-2^ Enzymes known as N-terminal acetyltransferases (NATs) carry out the modification by transferring an acetyl group from acetyl coenzyme A (CoA) to the N-terminal amino group of the substrate protein. Eukaryotic NAT family members – including NatA, NatB, NatC, NatD, and NatE – are known to function co-translationally, but post-translational N-terminal acetylation is observed for NatF, NatG, and possibly other NAT family members, including NatB.^3-5^ Unlike lysine side-chain acetylation, which is installed by lysine acetyltransferases and removed by lysine deacetylases,^6^ N-terminal acetylation is thought to be irreversible since no N-terminal deacetylase has been identified to date.

Notable among the members of the NAT family is NatB, which acts on ∼20% of the human proteins that are known to be N-terminally acetylated.^1^ NatB is selective for the N-terminus of proteins that retain the initiator methionine and have a Met-Asx/Glx (MD, ME, MN, or MQ) sequence as the first two residues. Such proteins are almost always N-terminally acetylated.^7^ As is the case with other members of its family, NatB regulates protein function and is therefore relevant in diseases, including cancer.^8-9^ N-terminal acetylation is important for α-synuclein (αS), which accumulates in aggregated deposits known as Lewy bodies (LBs) that are hallmarks of Parkinson’s disease (PD). N-terminal acetylation by NatB has been shown to affect αS aggregation *in vitro* and to influence levels of αS expressed in cellular models, both wild type (WT) and the disease mutant E_46_K.^10^ The same PTM is also thought to impact αS function in the synapse, altering its affinity for lipid vesicles^11-13^ as well as its interaction with protein partners.^14^ Recently, it has been shown that the acetylated N-terminus of αS plays an important role in the addition of new αS monomers to elongate amyloid fibrils.^15^ Therefore, there is significant interest in understanding the molecular basis for N-terminal acetylation of αS and the potential to modulate this to alter αS aggregation.

Human NatB (hNatB) consists of the catalytic subunit hNAA20 and the auxiliary subunit hNAA25, both of which are required for function.^16^ This subunit composition is conserved in yeast, in which deletion of either subunit gives rise to growth defects.^8^ A crystal structure of *Candida albicans* NatB (caNatB) has been reported with a peptide conjugate inhibitor,^17^ but hNAA20 and hNAA25 only share ∼40% and ∼20% sequence similarity with the corresponding caNatB subunit. Recently, we reported a cryo-electron microscopy (cryo-EM) structure of hNatB bound to an inhibitor consisting of CoA conjugated to a peptide of αS N-terminal residues 1-10, revealing structural differences with other NATs, as well as modes of recognition by hNatB specific to αS.^18^

The role that the remaining residues of αS play in interacting with NatB remains to be understood. Although NatB is known to be ribosome-associated and acetylates the nascent peptide, the enzyme also exists in a ribosome-unassociated form in the cytosol^3^. Given that some NATs have been found to have post-translational or even non-catalytic roles,^1^ it is possible that NatB also interacts with its substrates outside of its ribosome-bound state, with a chaperone-like function. In this work, we investigate the role of the αS central and C-terminal domains when engaging in a complex with NatB. We synthesize an inhibitor consisting of CoA covalently conjugated to full-length (FL) αS using native chemical ligation (NCL), demonstrating the first inhibitor of NatB consisting of a FL protein. Indeed, FL protein inhibitors of this type for any protein have rarely been made.^19-22^ Here, we take this type of semi-synthesis even further by combining NCL with unnatural amino acid (Uaa) mutagenesis,^23^ generating versions of the FL inhibitor with two orthogonally installed fluorescent probes for biophysical studies. We determine the cryo-EM structure of NatB bound to the CoA-αS_FL_ inhibitor. We then examine αS bound to NatB by single molecule Förster resonance energy transfer (smFRET) and reveal that binding induces conformational changes in αS, although it remains largely disordered. Computational simulations of the disordered regions of αS based on our NatB experimental cryo-EM structure help to explain the mechanism of conformational change and its implications for substrate recognition by NatB. These studies not only reveal important information about the function of NatB and its interactions with αS in particular, but also demonstrate efficient semi-synthetic strategies for triply functionalized protein-inhibitor fusions and how these can be combined with cryo-EM, smFRET, and computational modeling to address challenging structural biology targets.

## RESULTS

### Synthesis of Bisubstrate Inhibitor of NatB Consisting of Full-length αS

To synthesize a bisubstrate inhibitor of NatB consisting of CoA and FL αS, we performed NCL between a CoA-linked fragment of αS consisting of residues 1-10 and a partner fragment of consisting of residues 11-140 (Scheme 1). To generate the CoA-linked fragment αS_1-10_, we first carried out the synthesis of the peptide αS_1-10_ by fluorenylmethoxycarbonyl (Fmoc) based solid phase peptide synthesis (SPPS). To afford a C-terminal acyl hydrazide for ligation, 2-chlorotrityl resin was derivatized with Fmoc-hydrazide, and SPPS was carried out using standard conditions. After removal of the final Fmoc group from Met_1_ (**1**), bromoacetic acid was coupled to the N-terminus of the peptide as an active *O*-acylisourea using carbodiimide chemistry to afford **2**. Previous methods of bromoacetylation for CoA-linked inhibitor synthesis consisted of pre-activating a mixture of 10 equiv bromoacetic acid and 20 equiv *N,N’*-diisopropylcarbodiimide (DIC) in dichloromethane (CH_2_Cl_2_), removing the solvent *in vacuo*, and adding the mixture to the peptidyl resin in dimethylformamide (DMF).^24^ To maximize the yield of completely bromoacetylated product, we dissolved 10 equiv bromoacetic acid and 9.3 equiv DIC in DMF and added the mixture directly to the peptidyl-resin, a method employed in peptoid synthesis.^25^ Bromoacetylation proceeded to completion in 1 h. The bromoacetyl-αS_1-10_ acyl hydrazide **3** was obtained in 12% isolated yield.

To attach CoA via a nonhydrolyzable acetonyl linkage, the bromoacetyl-αS_1-10_ peptide **3** was reacted with CoA-SH. Whereas established bisubstrate synthesis methods react 1 equiv CoA-SH with peptide,^24^ we used excess CoA-SH to drive the reaction to completion. This was important because residual bromoacetyl-αS_1-10_ peptide leads to the formation of a side product resulting from nucleophilic attack of the thiol additive during ligation. Bromoacetyl-αS_1-10_ was redissolved in triethylammonium bicarbonate (TEAB) buffer pH 8, supplemented with 2 equiv CoA-SH, and the mixture was allowed to react for 4 h at room temperature and then overnight at 4 **°**C. After purification by reverse phase liquid chromatography (RP-HPLC), the CoA-linked αS_1-10_ peptide acyl hydrazide **4** was obtained in 76% isolated yield. The partner fragment, αS_11-140_ (**6**), was constructed by performing deletion polymerase chain reaction (PCR) on a plasmid of αS fused to a histidine-tagged intein (αS-Mxe-His_6_) and expressed in *E. coli* using the same method as previously described for αS C-terminal fragments.^26^ After thiol-mediated cleavage of the intein, methoxyamine-mediated deprotection of the N-terminal thiazolidine (Thz), and HPLC purification, the resulting αS_11-140_-C_11_ (**7**), bearing a Cys ligation handle, was obtained in an isolated yield of 20.9 mg per L of culture.

With fragments **4** and **7**, we performed NCL to build the bisubstrate inhibitor of NatB consisting of CoA-linked FL αS. CoA-αS_1-10_ acyl hydrazide **4** was dissolved in buffer at pH 3, chilled to -15 **°**C, and converted to an acyl azide using 10 equiv NaNO_2_. Following addition of 40 equiv mercaptophenyl acetic acid (MPAA) and 2 equiv αS_11-140_-C_11_ hydrazide **4** to generate MPAA thioester **5**, the reaction mixture pH was adjusted to 7.0, supplemented with reducing agent *tris*(2-carboxyethyl)phosphine (TCEP), and incubated at 37 **°**C with agitation. The reaction proceeded rapidly, with product (**8**) formation observed at 5 min and a moderate amount of product formation at 30 min (Figure 1). Despite the nucleophilicity of the lysine side chain, no side product was observed originating from the cyclization of the Lys_10_ thioester of αS_1-10_. After several hours, the ligation product was purified by HPLC and obtained in 68% yield. Finally, we converted the Cys_11_ ligation handle back to the native alanine by radical desulfurization. In the semi-synthesis of aggregation-prone proteins such as αS, it is strategic to minimize lyophilization of the full-length protein or aggregation-prone fragments. Alkyl thiol additives such as trifluoroethanethiol (TFET)^27^ or methyl thioglycolate (MTG)^28^ allow desulfurization in the same pot as ligation, eliminating a purification step. In this case, however, we found that the purified NCL product **8** was easily solubilized in buffer, likely because of the covalently linked CoA. We therefore took advantage of the faster-reacting MPAA-thioester, leading to rapid product formation and minimal hydrolysis of the reactive thioester. Thus, in a separate pot, the purified NCL product **8** was desulfurized using 20 mM radical initiator VA-044, 100 mM glutathione (GSH), and 250 mM TCEP. The resulting ligated product CoA-αS_FL_ (**9**) was purified by HPLC over a C4 column and obtained in 90% yield. Thus, we obtained the CoA-αS_FL_ bisubstrate inhibitor of NatB in 61% isolated yield overall. Note: A subsequent synthesis of **9** using ligation at residues 18/19 based on the route developed for labeled fluorescently labeled CoA-αS below produced protein in a slightly higher yield (see SI for details).

**Figure 1.**
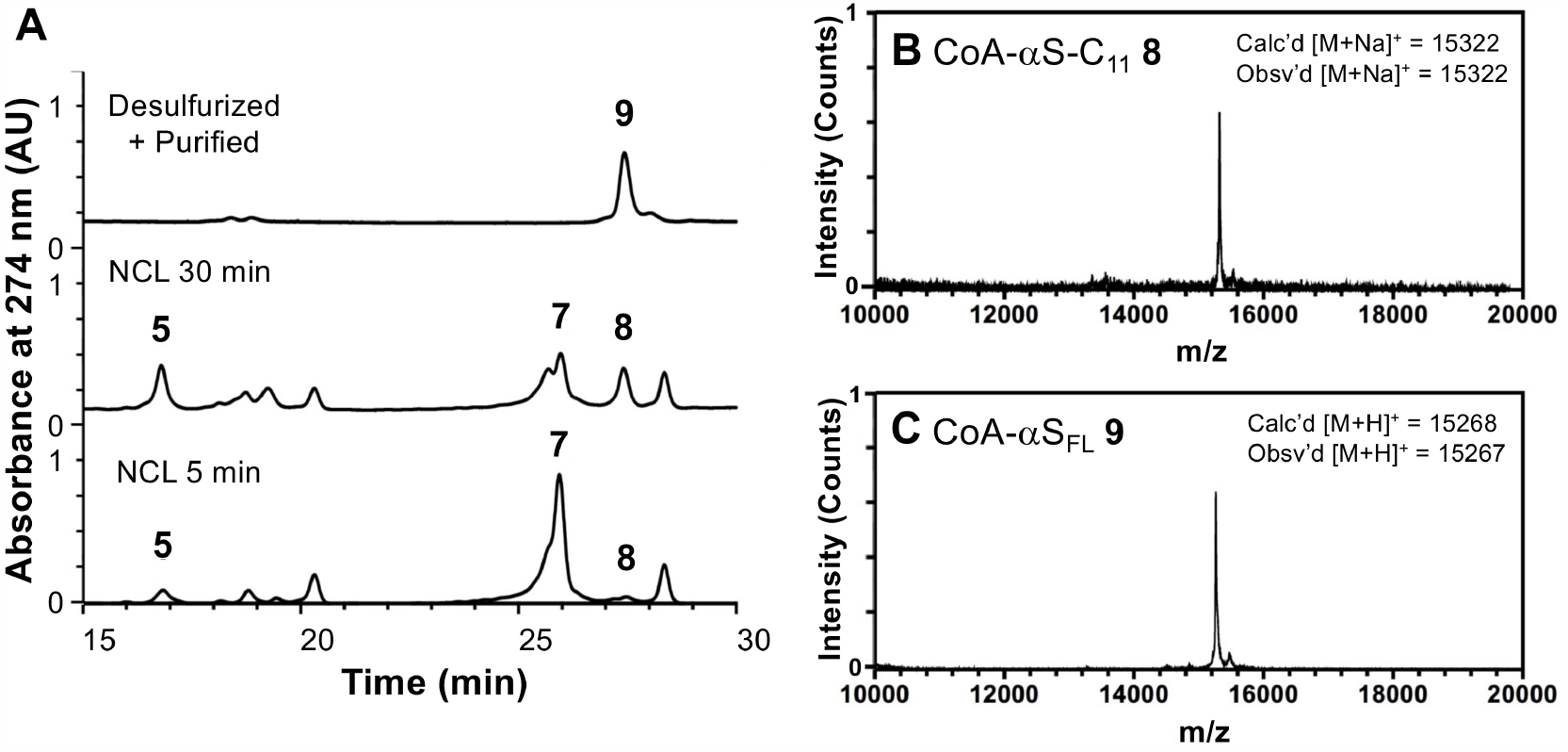
Semi-synthesis of CoA-αS_FL_ inhibitor. **(A)** Analytical HPLC (gradient: 10-50% B over 30 min) of NCL between CoA-αS_1-10_ MPAA thioester (**5**) and αS_11-140_-C_11_ (**7**) to afford the product CoA-αS_FL_-C_11_ (**8**) and desulfurized, purified product CoA-αS_FL_ (**9**). **(B)** MALDI-MS of ligation product CoA-αS_FL_-C_11_ (**8**). (**C**) MALDI-MS of desulfurized final product CoA-αS_FL_ (**9**).

### Potency of hNatB Inhibition by CoA-αS

We sought to compare the hNatB binding of our previously synthesized CoA-linked αS_1-10_ inhibitor^18^ to our novel inhibitor **9** consisting of FL αS. We found that the IC_50_ of the CoA-αS_1-10_ inhibitor was 1.63 ± 0.13 μM and the IC_50_ of FL inhibitor **9** was 2.19 ± 1.40 μM (Supporting Information, SI, Fig. S6). The comparable inhibition of hNatB by the CoA-αS_1-10_ and CoA-αS_FL_ inhibitors supports the conclusion that NatB preferentially acts on nascent polypeptides, with the residues beyond the first 10 not contributing significantly to the affinity of the enzyme for the substrate. However, we were still interested in how binding to hNatB influenced the conformation of αS, which could have important implications for its aggregation, even though this interaction is thought to occur exclusively or predominantly co-translationally.

### Structure of hNatB in Complex with CoA-αS_FL_ by Cryo-EM

We determined the structure of the CoA-αS_FL_ inhibitor **9** bound to hNatB by single particle cryo-EM (Fig. 2). We found that the first 5 residues of αS were engaged in clear interactions with hNatB, whereas the residues after αS Met_5_ were not resolved (Fig. 2), as previously seen with the peptide-based inhibitor.^18^ Despite the presence of full length human αS, no density was observed that could be unambiguously assigned to αS for residues 6-140. (Fig. 2 and SI, Fig. S7). The fact that no extra residues were resolved suggests that the conformational heterogeneity of the rest of αS is maintained in the complex, showing that αS remains intrinsically disordered even in complex with hNatB. Thus, we turned to methods more suited to disordered protein to determine whether the conformations of the other αS domains were altered by hNatB binding.

**Figure 2.**
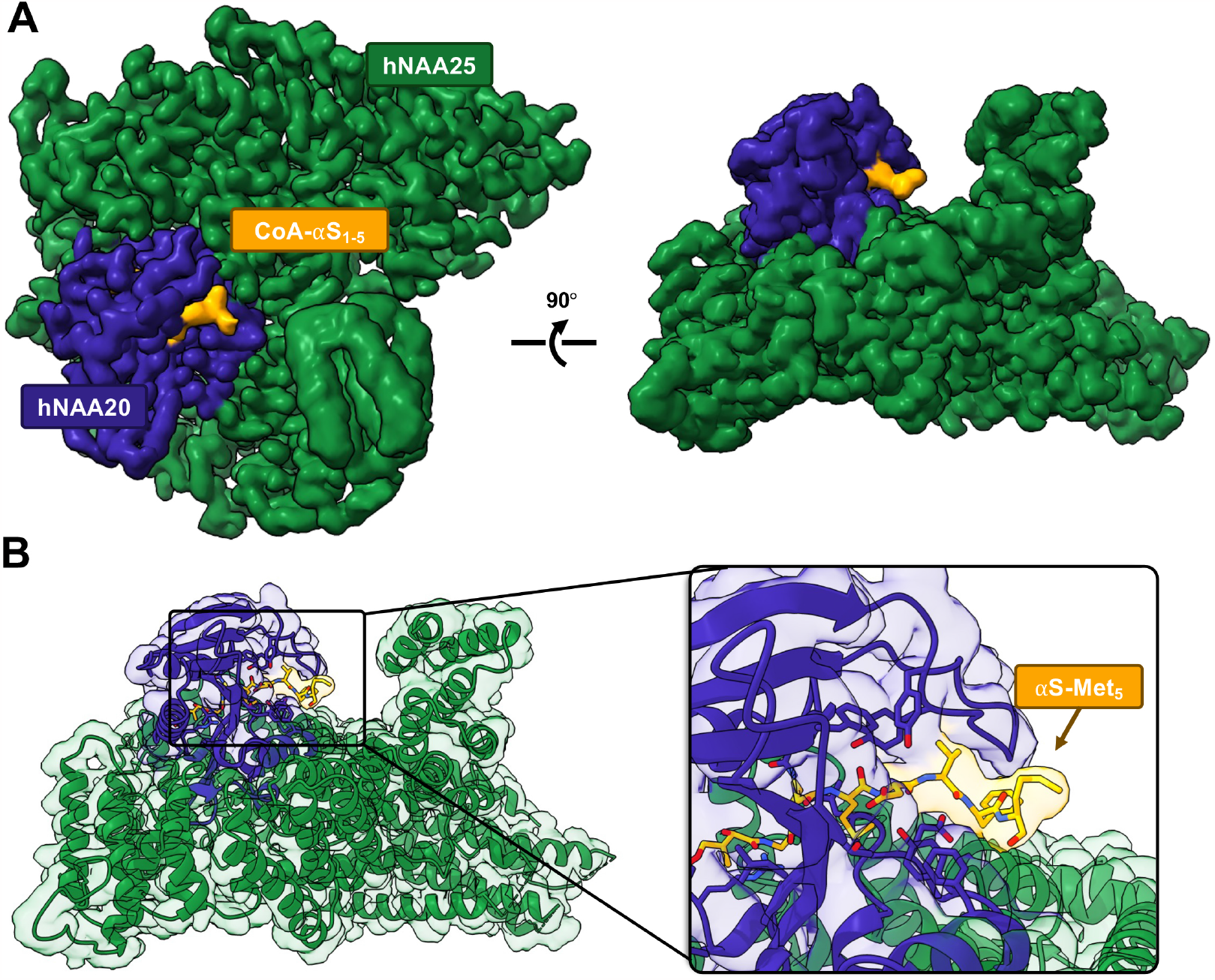
Cryo-EM structure of hNatB in complex with CoA-αS_FL_ inhibitor. (**A**) The cryo-EM map rendered in Chimera with subunits hNAA25 (green) and hNAA20 (purple) and CoA-αS_FL_ (yellow). (**B**) The cryo-EM model of hNatB and CoA-αS_FL_ (colored as in A). Zoomed in view highlights bound CoA-αS_FL_. There is no interpretable cryo-EM map density beyond αS residue Met_5_.

### Synthesis of Doubly Fluorescently Labeled Bisubstrate Inhibitor CoA-αS_FL_

To assess changes in the conformational ensemble of the regions of αS beyond the hNatB-bound N-terminus, we turned to smFRET. For these studies, we needed to synthesize versions of αS not only fused to CoA, but also bearing two fluorophores. We recognized that we could attach one fluorophore via the Cys residue used for NCL. For the other fluorophore, we aimed to replace fragment **7** used above with a C-terminal αS fragment carrying the alkynyl amino acid propargyl tyrosine (π), incorporated by amber codon suppression,^29^ for attaching a fluorophore via click chemistry. Preliminary attempts to ligate at position 10 were hindered by low expression yields of the αS_11-140_-C_11_π_n_ (n=72, 94, 136) fragments as compared to the all-canonical αS_11-140_-C_11_ (**7**) fragment, so we sought to maximize the yield of the NCL product by choosing a more optimal ligation site: one that can afford a more rapid thioester exchange with little possibility for side reactions. To take advantage of a sterically unencumbered Ala_17_/Ala_18_ ligation site, we synthesized N-terminal fragment CoA-αS_1-17_ and attempted to express the thiazolidine-protected αS_18-140_-C_18_π_n_ (n=72, 94, 136; **S3a, S3b, S3c**, respectively) constructs. We expected, as with other αS C-terminal fragments, that after initiator methionine cleavage by methionine aminopeptidase (MAP), reaction of the exposed N-terminal cysteine with various carbonyls in the cell would afford the desired product as Thz-protected derivatives. However, we observed near-exclusive production of a fragment missing the Cys_18_ ligation handle that we had attempted to introduce and only trace amounts of the desired fragment (SI, Fig. S8). The excess cleavage was possibly due to an unexpected activity of MAP (see Discussion). We therefore switched our semi-synthesis strategy to ligate at Ala_18_/Ala_19_ instead (Scheme 2), allowing us to still take advantage of a rapidly ligating alanine thioester. We synthesized CoA-αS_1-18_ acyl hydrazide (**S2**) and obtained the product in 26% isolated yield after SPPS and reaction of the bromoacetyl peptide with CoA-SH. The C-terminal fragments were expressed with propargyl tyrosine at position 72 (**11a**), 94 (**11b**), or 136 (**11c**) to generate inhibitors labeled at different positions of the protein, allowing probing of domain-specific changes in conformation by smFRET: Residue 72 is in the center of the hydrophobic, aggregation-driving region; residue 94 is at the end of the membrane-binding domain; and residue 136 is near the C-terminus. After Thz-deprotection, 7.7 mg of pure αS_19-140_-C_19_π_72_ (**11a**), 15.7 mg of αS_19-140_-C_19_π_94_ (**11b**), and 8.5 mg of αS_19-140_-C_19_π_136_ (**11c**) were obtained per L of bacterial culture. NCL reactions between these fragments and CoA-αS_1-19_ MPAA thioester (**10**, generated *in situ* using the procedure described for conversion of **4** to **5**) were carried out as for the non-fluorescent FL inhibitor except using 1.2 equiv C-terminal partner (Fig. 3). The ligation product CoA-αS-C_19_π_72_ (**12a**) was obtained in 25% yield, CoA-αS-C_19_π_94_ (**11b**) in 30% yield, and CoA-αS-C_19_π_136_ (**12c**) in 23% yield. Instead of desulfurization, the Cys_19_ of the ligation product was labeled with Alexa Fluor 488 (AF488) maleimide. Following purification, the singly labeled products CoA-αS-C^488^_19_π_72_, CoA-αS-C^488^_19_π_94_, and CoA-αS-C^488^_19_π_136_ were each click labeled using Alexa Fluor 594 (AF594) azide and a catalytic mixture consisting of CuSO_4_, tris-hydroxypropyltriazolylmethylamine (THPTA) ligand, and sodium ascorbate. Labeling reactions were quantitative. The final products, labeled with two fluorophores and the CoA bisubstrate inhibitor, CoA-αS-C^488^_19_π^594^_72_ (**13a**), CoA-αS-C^488^_19_π^594^_94_ (**13b**), and CoA-αS-C^488^_19_π^594^_136_ (**13c**) were purified by HPLC for use in fluorescent studies. As noted above, the 18/19 ligation site was used for a subsequent synthesis of CoA-αS_FL_ (**9**).

**Figure 3.**
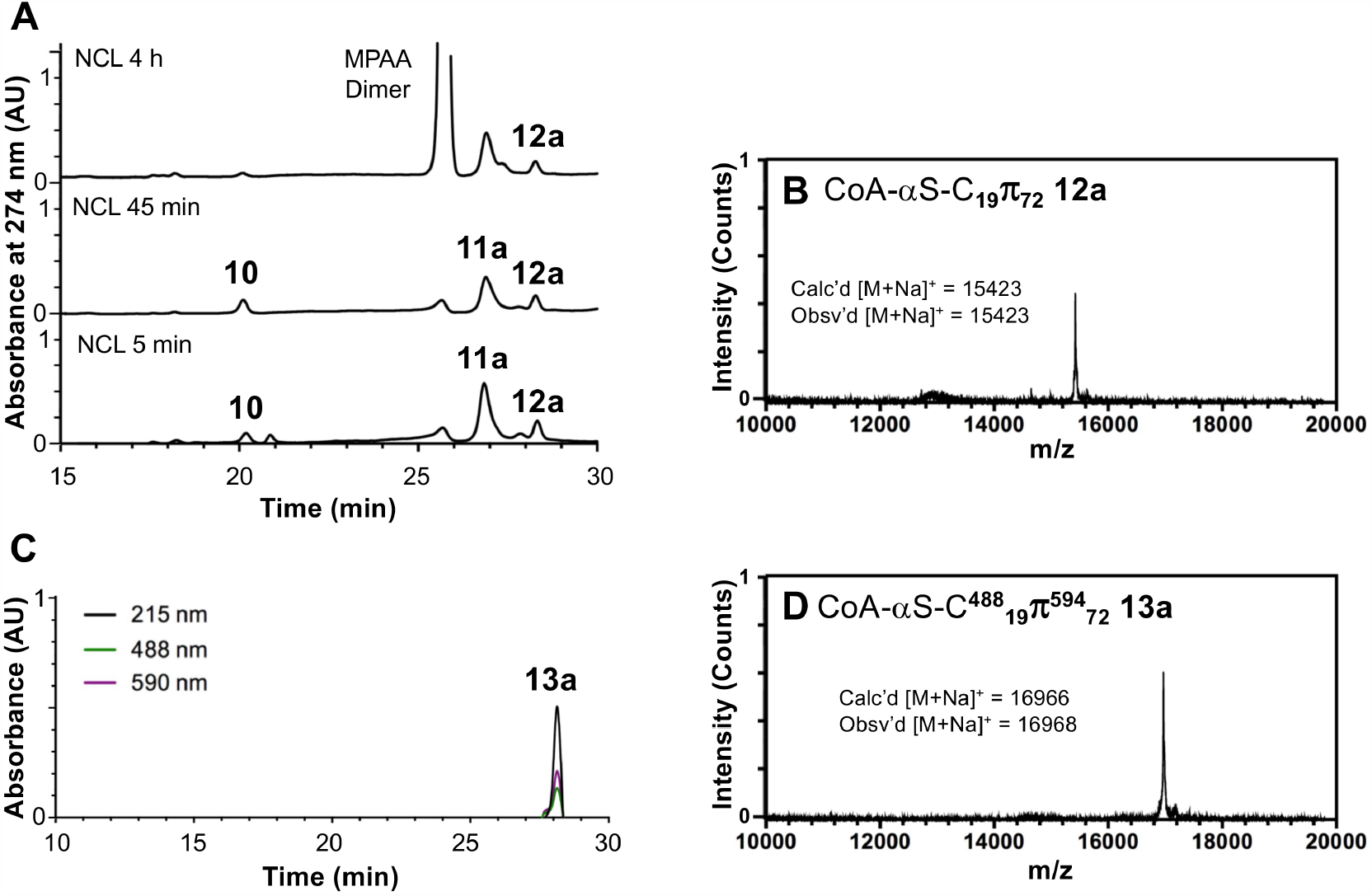
Semi-synthesis of doubly fluorescent CoA-αS_FL_ inhibitor. (**A**) Analytical HPLC of NCL between CoA-αS_1-18_ MPAA thioester (**10**) and αS_19-140_-C_19_ π_72_ (**11a**) to afford the product αS_19-140_-C_19_π_72_ (**12a**). The MPAA Dimer peak is intentionally cutoff as this additive is present in high abundance. (**B**) MALDI-MS of ligation product αS_19-140_-C_19_π_72_ (**12a**). (**C**) Analytical HPLC of AF488-maleimide and AF594-N_3_ labeled and purified product CoA-αS-C^488^_19_π^594^_72_ (**13a**) (Retention time 28.0 min). (**D**) MALDI-MS of CoA-αS-C^488^_19_π^594^_72_ (**13a**). HPLC gradients 10-50% B over 30 min.

### Conformation of CoA-αS_FL_ in the Presence of hNatB by smFRET

To ensure the homogeneity of the enzyme-inhibitor complex to be examined by smFRET, we determined the molar equivalents of hNatB needed for full binding of CoA-αS_FL_. Whereas preparation of the complex for structural studies involved the use of excess inhibitor, smFRET measurements of the hNatB/CoA-αS complex are simplified by the absence of free inhibitor in solution so that all of the fluorescence derives from the bound species. We determined the molar equivalents of hNatB necessary for full complex formation by fluorescence correlation spectroscopy (FCS), a technique in which fluctuations in fluorescence intensity from a fluorescently labeled sample are temporally autocorrelated to deduce physical information, including concentration and diffusion time. For our purposes, the autocorrelation curves were fit to a function describing a single diffusing species to determine the diffusion time of the fluorescently labeled inhibitor CoA-αS-C^488^_19_π_n_ (n = 72, 94, 136) free in solution (see SI Methods for details of FCS data fitting). We then formed enzyme-inhibitor complexes by incubating CoA-αS-C^488^_19_π_n_ with various molar equivalents of hNatB: 1, 10, 40, or 100 equiv. The diffusion time of each complex was determined by fitting these autocorrelation curves to the same single-component equation as for the free inhibitor. Saturation of inhibitor binding was confirmed as the diffusion time reached a plateau at 40 molar equivalents of hNatB. Note that the requirement of a large excess of hNatB is expected because the FCS and smFRET experiments are done at CoA-αS concentrations well below the K_D_ for the hNatB/CoA-αS complex, expected to be in the μM range based on its IC_50_.

To study the conformational changes of CoA-αS_FL_ in the presence of hNatB, the energy transfer efficiency (ET_eff_) between donor and acceptor fluorophores was measured for the three pairs of labeling positions on the FL inhibitor. CoA-αS labeled with AF488 and AF594 (**13a, 13b**, or **13c**) was measured in buffer and compared with the inhibitor in complex with the enzyme (Fig. 4). The complex was pre-formed by incubating CoA-αS with 100 equiv hNatB, followed by dilution into the measurement chamber. We found that the first half of CoA-αS displayed no significant change in conformation upon hNatB binding, as ET_eff_ between positions 19 and 72 remained the same for free and bound inhibitor. On the other hand, ET_eff_ between positions 19 and 94 decreased from 0.48 to 0.43 upon binding to hNatB, indicating an expansion between the N-terminus and a distal portion of CoA-αS. Similarly, ET_eff_ between positions 19 and 136 decreased from 0.30 to 0.25 upon binding, showing an expansion of the N- and C-termini of CoA-αS. Based on these ET_eff_, the root mean square (RMS) distance were inferred using a Gaussian chain polymer model.^30-31^ The RMS distance between positions 19 and 94 increased from 64 Å in solution to 68 Å when NatB-bound. Although a similar decrease in ET_eff_ was seen between positions 19 and 136, these lower efficiencies corresponded to a larger increase in RMS distance, from 83 Å to 91 Å, as ET_eff_ scales inversely with the sixth power of the distance between the fluorophores. For all pairs of positions examined, a broadening of ET_eff_ distribution was observed, possibly indicative of increased conformational sampling or slower dynamics.

**Figure 4.**
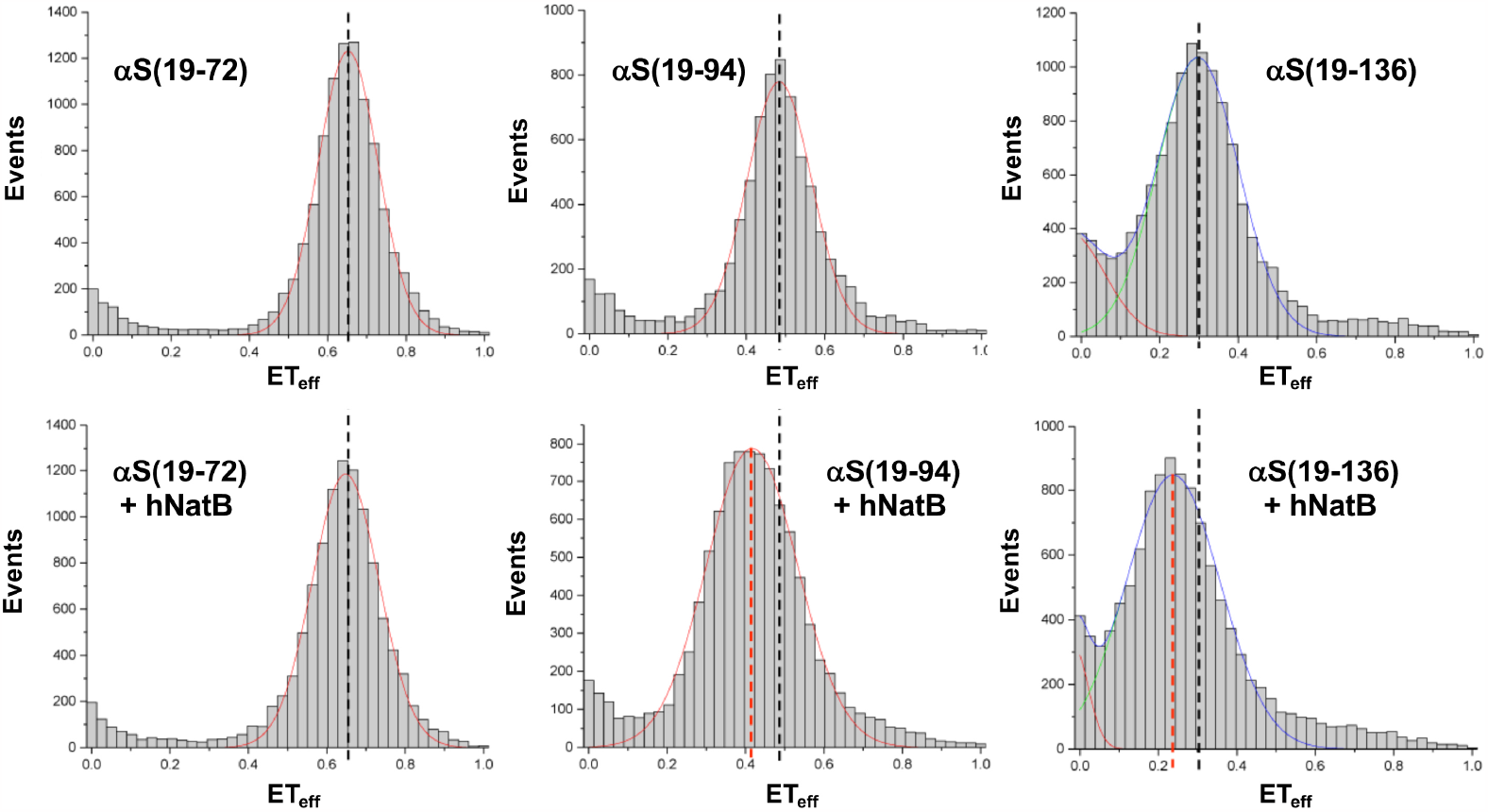
Conformation of CoA-αS_FL_ in complex with hNatB by smFRET. Representative single molecule FRET histograms fit to Gaussian distributions. Mean FRET efficiencies (ET_eff_) are indicated for CoA-αS_FL_ alone (black dashed lines) and in complex with hNatB (red dashed lines). The histograms show no change in ET_eff_ between residues 9 and 72 upon complex formation with hNatB; change in ET_eff_ consistent with expansion between positions 9 and 94 as well as between positions 9 and 136. αS(19-n) refers to measurements made with CoA-αS-C^488^_19_π^594^_n_ (**13a, 13b**, or **13c**). The complete set of histograms and details of data fitting are given in the SI.

### Rotational Mobility of CoA-αS_FL_ Domains in Complex with hNatB

We turned to fluorescence anisotropy measurements to gain further insight into whether the interaction between CoA-αS and hNatB involved the folding back of distal residues of αS onto the enzyme. We determined the fluorescence anisotropy of each labeled CoA-αS inhibitor, which provides information on the rotational mobility at that residue position. We and others have previously shown that such anisotropy (or polarization) measurements can be valuable probes of local conformational flexibility within αS.^32-33^ By comparing the anisotropy in the presence and absence of hNatB, we could ascertain whether distal residues of the inhibitor were interacting with the enzyme, as restrained rotation of the polypeptide portion of interest as a result of binding would lead to a large increase in anisotropy. In the absence of hNatB, CoA-αS labeled with AF594 was characterized by anisotropy values of 0.12 ± 0.01 for position 72, 0.11 ± 0.01 for position 94, and 0.11 ± 0.01 for position 136 (SI, Fig. S26). In complex with hNatB, a slight increase in anisotropy, by ≤ 1.5-fold, was observed for all positions of the inhibitor. The extent of the increase was expected based on slower tumbling of the complex as a result of inhibitor binding to hNatB, but was not large enough in magnitude to indicate a restriction of rotational mobility of the CoA-αS residues. A previous report examining lipid interactions of αS by anisotropy took advantage of tryptophan fluorescence using site-specific mutants, and the researchers observed a 3-fold increase in anisotropy for the αS N-terminus, the helical transition of which imparts rotational restriction, but a < 2-fold increase for the C-terminal domain, which experiences no restriction to rotation upon binding to lipid vesicles.^32^ The small increase in anisotropy of our CoA-αS inhibitor upon binding to hNatB implies very little inhibition of rotational dynamics of distal domains, implying that the conformational dynamics of the αS C-terminus are not affected by the presence of the enzyme.

### Structural Modeling of αS Disordered Regions in hNatB Complex

To better understand the conformational ensemble of the disordered regions of αS while in complex with hNatB, we simulated the complex in PyRosetta using the cryo-EM structure as a starting point. Input structures were processed according to Ferrie *et al*. for FastFloppyTail (FFT).^34^ hNatB was fixed during course grain sampling of the αS structure, and side chains were allowed to move during full atom refinement. The 1000 lowest scoring structures were relaxed and used in generating quantitative structural data for this unrefined ensemble, which we refer to as the raw ensemble. From these structures, the average radius of gyration (R_g_) for αS was 44.3 Å, which is an increase of 13.3 Å from simulations of free αS using FFT, as well as a 17.7 Å increase from the free αS NMR derived R_g_ of 26.6 Å.^35^ This R_g_ value is consistent with an extended structure based on smFRET observations. Specific comparisons of the distance distributions derived from the smFRET measurements to C_α_-C_α_ distances from the raw simulated ensemble are shown in SI (Fig. S30). The changes in inter-residue distances between αS alone and αS in complex with hNatB are shown in Figure S27, highlighting the extension of the C-terminal region with inter-residue distance increasing by as much as 40 Å (the individual contact map for simulated αS/hNatB is shown in Fig. S27, SI). These simulations indicate that it is unlikely that the increase in C-terminal smFRET distances originates from specific interactions of αS with hNatB, with no region consistently contacting hNatB. Outside the binding pocket, the most significant area of interaction involves residues 13-20 of hNAA25 contacting residues 15-20 of αS, which occurs in 20% of the structures in the raw ensemble (determined with a 10 Å cut-off and occurrence in at least 5% of the ensemble SI, Fig. S30). These interactions are transient in nature, but may contribute to conformational changes resulting in the observed changes in smFRET. Sequestration of the N-terminus would reduce favorable charge contacts with the C-terminus, leading to an overall expansion of the protein.

While we were pleased to see that unconstrained simulations of the αS/hNatB complex were consistent with the experimental smFRET and FCS data, we wished to observe whether refinement of the ensemble using the smFRET data as constraints could provide additional insight into αS/hNatB interactions. To achieve our objective, we added an additional score term based on the average distance derived from the smFRET measurement as well as the breadth of the distribution for each of the three residue pairs (for rescoring details, refer to the SI, and for reweighted scores of the refined ensemble, see Figure S28). By utilizing this constraint, we generated a refined ensemble that more accurately reflects the distribution of distances observed in smFRET measurements (Fig. S29) and comprises 262 members. The 10 lowest energy structures from the refined ensemble are shown in Figure 5A. The average R_g_ for the refined ensemble (Fig. 5B) was identical to that of the raw ensemble (44.5 Å and 44.3Å, respectively), with only minor shifts in populations, indicating that refinement did not significantly skew the ensemble (SI Fig. S27 and S28). Comparison of the inter-residue distances from αS residues 20-100 in the hNatB refined ensemble to those in the free αS ensemble shows that significant compaction occurs in the presence of hNatB (Fig. 5C). More compaction in these regions is observed in the refined ensemble than in the raw ensemble (SI, Fig. S27 and S29), as expected in the presence of the 19-72 smFRET distance constraint. As a consequence, more transient interactions are observed for refined ensemble. However, αS contact with residues 13-20 of hNAA25 remains the only consistent interaction in the refined ensemble, occurring in 20% of the structures (Fig. 5 and SI, Fig. S30). Taken together, the simulations support the observations from cryo-EM, smFRET, and anisotropy that the unbound portion of αS remains largely disordered in the presence of hNatB, but that a weak interaction occurs in this region.

**Figure 5.**
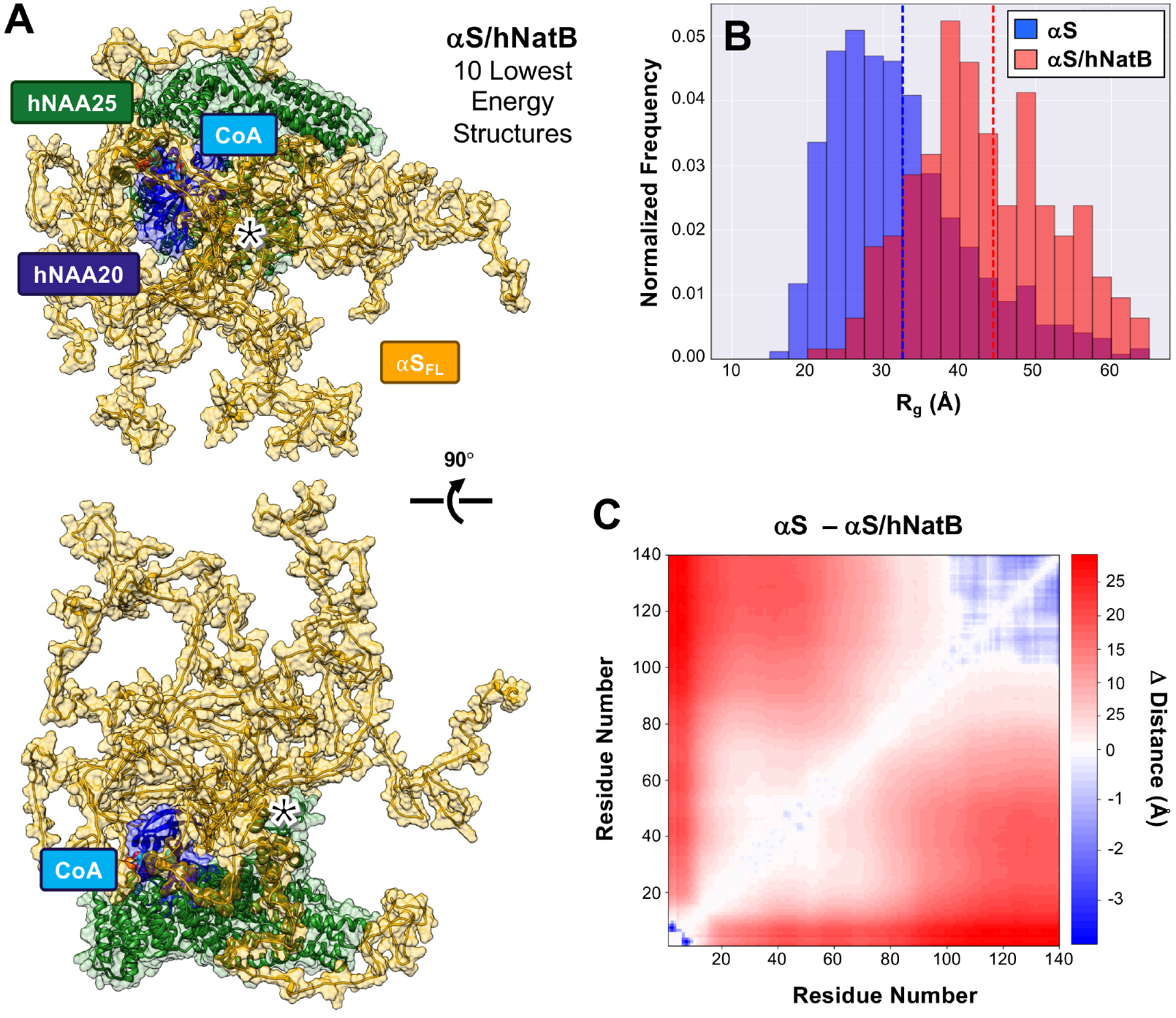
Simulations of CoA-αS_FL_ in complex with hNatB. (**A**) The CoA-αS_FL_/hNatB complex was simulated in PyRosetta, using the cryo-EM structure as a starting point. The resulting ensemble was then refined using smFRET-derived distance constraints. The 10 lowest energy structures of the αS/hNatB complex from the refined ensemble are shown in the same two orientations as in Figure 2, with transparent space-filling models superimposed in both images. The 13-20 region of hNAA25, which contacts αS residues 15-20 in 20% of the ensemble, is denoted with a *. (**B**) The radius of gyration (R_g_) of the 1000 lowest energy structures in the free αS (blue) or refined αS/hNatB (red) ensembles. Dashed lines indicating the ensemble averages are colored accordingly. (**C**) A difference heat map showing the changes in the average inter-residue distances between the αS and refined αS/hNatB ensembles.

## DISCUSSION

αS is subject to numerous PTMs, and among them N-terminal acetylation stands out, as αS is constitutively acetylated in living organisms. The modification of αS by NatB has important implications for proper protein function and disease. We recently determined the modes of substrate recognition by hNatB specific to αS.^18^ In the prior study, we revealed that hNatB accommodates Met_1_ of αS in a hydrophobic pocket and forms hydrogen bonds with Asp_2_ of αS mainly using a key histidine residue (hNatB-His_73_) in the substrate-binding site, consistent with the known substrate profile of NatB for proteins with a M-D/E/N/Q as starting residues. The third and fourth residues of αS engage less extensively with hNatB residues, and the first ∼5 residues of a CoA-αS_1-10_ inhibitor are clearly resolved in the cryo-EM structure of the complex with hNatB. In this study, we found that an inhibitor consisting of FL αS also displays clearly resolved structure in the first ∼5 residues when bound to hNatB with the same interactions observed previously (Fig. 2), but that the rest of the protein is unresolved in the cryo-EM map, presumably due to structural heterogeneity.

To examine the conformational ensemble adopted by distal domains of αS, we turned to smFRET using fluorescently labeled CoA-αS_FL_ at various pairs of positions. As the synthesis of a full-length protein conjugate with multiple modifications can be challenging, we sought multiple routes to synthesize CoA-αS_FL_. We optimized the semi-synthesis of the unlabeled CoA-αS_FL_ for structural characterization and further modified our route to the doubly fluorescently labeled inhibitor for single molecule studies. Some considerations of note are as follows. Since the yield of protein expression for the αS C-terminal fragment is lower for the unnatural amino acid variant, we switched the ligation site to the more optimal 18/19 position. Upon re-synthesis of CoA-αS_FL_, we found that the 18/19 site was also superior for this ligation. To have more of the C-terminal fragment available for NCL (which is optimal at mM concentrations), we carried out the ligation first, followed by maleimide labeling of Cys_18_ and click chemistry labeling of the π residue. In the earlier iteration with ligation at position 10, we click labeled first in order to take advantage of our previously observed deprotection of the Thz residue under the Cu-catalyzed click conditions,^36^ but found that this limited the amounts of protein available for the ligation step.

In our semi-synthesis of the FL inhibitor using positions 17/18 in αS as the ligation site, the loss of Cys_18_ from Thz-αS_18-140_-C_18_π was possibly due to the activity of MAP in *E. coli*. Although MAP is known to cleave the initiator Met when the next amino acid (at position P1’) is small, systematic variation of amino acids at positions P2’ to P5’ in substrate peptides has demonstrated that the identity of the P2’ residue has some effect on enzyme activity.^37^ More specifically, a triple mutant of MAP, engineered to be more efficient, has been demonstrated to remove the P1’ residue when the antepenultimate residue P2’ is also small.^38^ It is possible that WT MAP may do the same, especially if MAP can recognize as a substrate not only the linear thioether in methionine, but also the cyclic thioether in Thz-protected cysteine. In our case, the Cys_18_ at P1’ may have been removed due to the small size of Ala_19_ at P2’. It is also possible that Cys_18_ was removed by aminopeptidases other than MAP. It has been observed previously, albeit specifically for a proline at P2’, that both the initiator methionine and an alanine at P1’ are retained when incubated with MAP *in vitro*, despite being removed in *E. coli* hyperproducing MAP.^39^

Nevertheless, we successfully synthesized fluorescently labeled CoA-αS_FL_ by performing NCL at positions 18/19 instead, taking advantage of the rapidly reacting Ala_18_ thioester and introducing Cys_19_ as a ligation handle. We also made use of the ligation handle to attach the AF488 fluorophore with thiol-maleimide chemistry and orthogonally introduced the AF594 fluorophore through copper-catalyzed azide-alkyne cyclization.

Prior NMR and FRET studies have found evidence of long-range favorable electrostatic interactions between the termini of αS in solution.^40-44^ In this study, our smFRET measurements show an increase in the distance between the N-terminus (residue 19) and the end of the membrane binding domain (residue 94) as well as between the N- and C-termini (residue 136) of αS when bound to hNatB, reflecting diminishment of those interactions. Neither our cryo-EM structure nor our anisotropy measurements find evidence for stable interactions between αS and hNatB beyond the N-terminal residues involved in binding. We do not observe any changes in the N-terminal half of the protein (residues 19 and 72) upon hNatB binding by smFRET. However, as the last 40 residues bear a -13 charge, it may not be surprising that this region is more sensitive to the charge screening effects of hNatB. The FRET-based RMS distances between positions 19 and 94 increased from 64 Å in solution to 68 Å when NatB-bound, while for positions 19 and 136, the increase in RMS distance was from 83 Å to 91 Å. For both of these constructs, the dimensions of αS in the NatB bound state are consistent with a very extended backbone, more so than expected for a random coil.

The structures in the raw simulations of αS bound to hNatB are consistent with the smFRET data. For the specific residue pairs probed by smFRET, the changes in distance trend with the experimental measurements. While the simulated distance distributions are broader than those seen in the smFRET data, this may in part be a consequence of differences in the sampling regimes in the experiment versus the PyRosetta Monte Carlo simulations^45^ or in insufficient inclusion of broadening effects in the smFRET distance conversion.^46^ Refinement of the ensemble resulted in distance distributions that were more consistent with smFRET data, without changing the overall character of the αS ensemble. As noted above, the extension of the C-terminus does not seem to be the result of major interactions with hNatB, where only the 13-20 region of hNAA25 demonstrated any significant contact propensity with αS in either ensemble. Since this is a lower resolution region of the cryo-EM structure (SI, Fig. S7) and it interacts with the site at which αS was fluorescently labeled, additional studies will be necessary to confirm the relevance of the computational finding. When considered as a whole, our data are consistent with a model in which favorable electrostatic interactions in αS are screened by the sequestration of the positively charged N-terminal residues of αS within NatB, resulting in a more extended C-terminal region of the protein. This implies that if soluble NatB were to interact with αS away from the ribosome, no stable complex would be formed. However, since breaking of the N- and C-terminal contacts in αS has been shown to make αS more aggregation-prone,^41^ this type of interaction is unlikely to contribute to a chaperone-like activity for NatB. It may actually have the opposite effect, implying that a reduction in NatB interactions could be beneficial. Finally, it is interesting to consider that the absence of any stable interactions with client proteins beyond the N-terminus may be a useful attribute for NatB, where it must act transiently on the co-translational complex and then dissociate to act on another nascent protein.

## CONCLUSION

NatB is known to associate with its substrates when they are tethered to the ribosome, and residues beyond the first few that were resolved in our cryo-EM structures could also potentially play a role in substrate recognition. For NatA, it has been shown that 40-100 amino acids of a substrate need to be present for acetylation to take place, but this may be simply to clear the exit tunnel for enzyme access.^47^ In the case of hNatB studied here, the distal residues of αS do not seem to significantly contribute to specificity, as seen from the comparable IC_50_ values of the CoA-αS_1-10_ and CoA-αS_FL_ inhibitors and from the lack of additional density attributable to αS in the CoA-αS_FL_/hNatB cryo-EM data. While there are no stable interactions between hNatB and αS beyond the first few residues, the conformation of the αS_FL_ polymer is altered by the presence of the enzyme. Our smFRET, anisotropy, and modeling studies indicate that sequestering the N-terminus of αS in the hNatB active sites releases the C-terminus. For ribosomally-bound NatB, the absence of stable interactions beyond the first few residues may have been evolutionarily tailored, as this would prevent it from interfering with protein translation and allow NatB to easily dissociate to engage another client protein. If NatB-substrate interactions can take place freely in the cytosol (unassociated yeast NatB subunits have been found in the cytosol),^3^ the induced conformational changes seen upon αS binding may lead to effects on protein folding, as the breaking of N- and C-terminal interactions has been shown to increase the aggregation propensity of αS.^40-41^ Finally, the relevance of hNatB to many diseases makes it an important biological target: A recent report has pointed to inhibition of hNatB as a therapeutic strategy against synucleinopathies,^10^ highlighting the importance of our study of the behavior of a FL αS-conjugate inhibitor in the presence of hNatB. While the αS/hNAA25_303-310_ interaction only occurs in 20% of structures, further investigation is warranted, as targeting this region points to a potential mechanism for inhibiting interactions of αS with hNatB. Targeting this site, rather than the active site where CoA and the peptide substrate N-terminus bind, could allow one to specifically inhibit αS interactions with hNatB, avoiding undesired side effects of general hNatB inhibition.

Our ability to semi-synthesize an enzyme inhibitor consisting of a co-substrate-linked to a full-length protein has broad implications. In a review of the literature, we have identified only four prior examples of the semi-syntheses of protein probes to trap an enzyme in a conformation of interest.^22, 48^ A ubiquitin- (Ub-) adenylate mimic has been used to trap Ub E1 ligase in an open conformation before pyrophosphate release, a Ub-vinylsulfonamide has been semi-synthesized to covalently attach to the E1 catalytic cysteine, trapping the enzyme in a closed conformation, an ATP conjugate of protein kinase A has been used to study its mechanism, and a difluoromethylphosphonate analog of phosphothreonine has been used in dissecting the mechanism of protein phosphatase-1 autoinhibition.^19-21, 48^ Our semi-syntheses not only functionalize a full-length protein with a covalently linked chemical group for enzyme trapping and structural biology, but also orthogonally incorporate two fluorescent probes for single molecule spectroscopy study of conformational dynamics that are not accessible to cryo-EM. The cryo-EM and smFRET data obtained with these constructs then enable computational models of the full protein complex, including the disordered regions. Our successful semi-syntheses demonstrate that analogous inhibitors can be used to capture other challenging protein targets, facilitating future studies of enzyme structure and dynamic protein substrate interactions.

## ASSOCIATED CONTENT

### Supporting Information

Detailed protocols for cryo-EM, single molecule FRET, fluorescence correlation spectroscopy, peptide synthesis, protein expression, ligation and purification, and computational modeling as well as additional characterization and data (PDF) are available free of charge at http://pubs.acs.org.

## AUTHOR INFORMATION

### Author Contributions

B.P., S.G, K.Sc., S.D., E.R., R.M., and E.J.P designed the experiments. R.P. and E.J.P. designed the computational simulations, and they were performed by R.P. B.P., S.G., K.Sc., S.D., K.Sa., and M.S. performed the experiments and data analyses. The manuscript was written with input from all authors. All authors have given approval to the final version of the manuscript.

## Acknowledgments

This research was supported by the National Institutes of Health (NIH R01 NS103873 to E.J.P., R01 NS120625 to E.R., R35 GM1118090 to R.M.) Instruments supported by the National Science Foundation and NIH include matrix-assisted laser desorption ionization mass spectrometers (NSF MRI-0820996, NIH S10 OD030460). B.P. thanks the University of Pennsylvania for support through a Dissertation Completion Fellowship. R.P. and K.Sc. were supported by the NIH Chemistry Biology Interface Training Program (T32-GM133398). S.M.G. was supported by the NIH Structural Biology and Molecular Biophysics Training Program (T32-GM132039). M.S. thanks the Nakajima Foundation for scholarship funding. K.Sa. thanks the funding from Shizuoka University for fostering joint international research. R.P. and E.J.P. thank Sam Giannakoulias for helpful discussions of computational modeling.

---

**Scheme 1.**
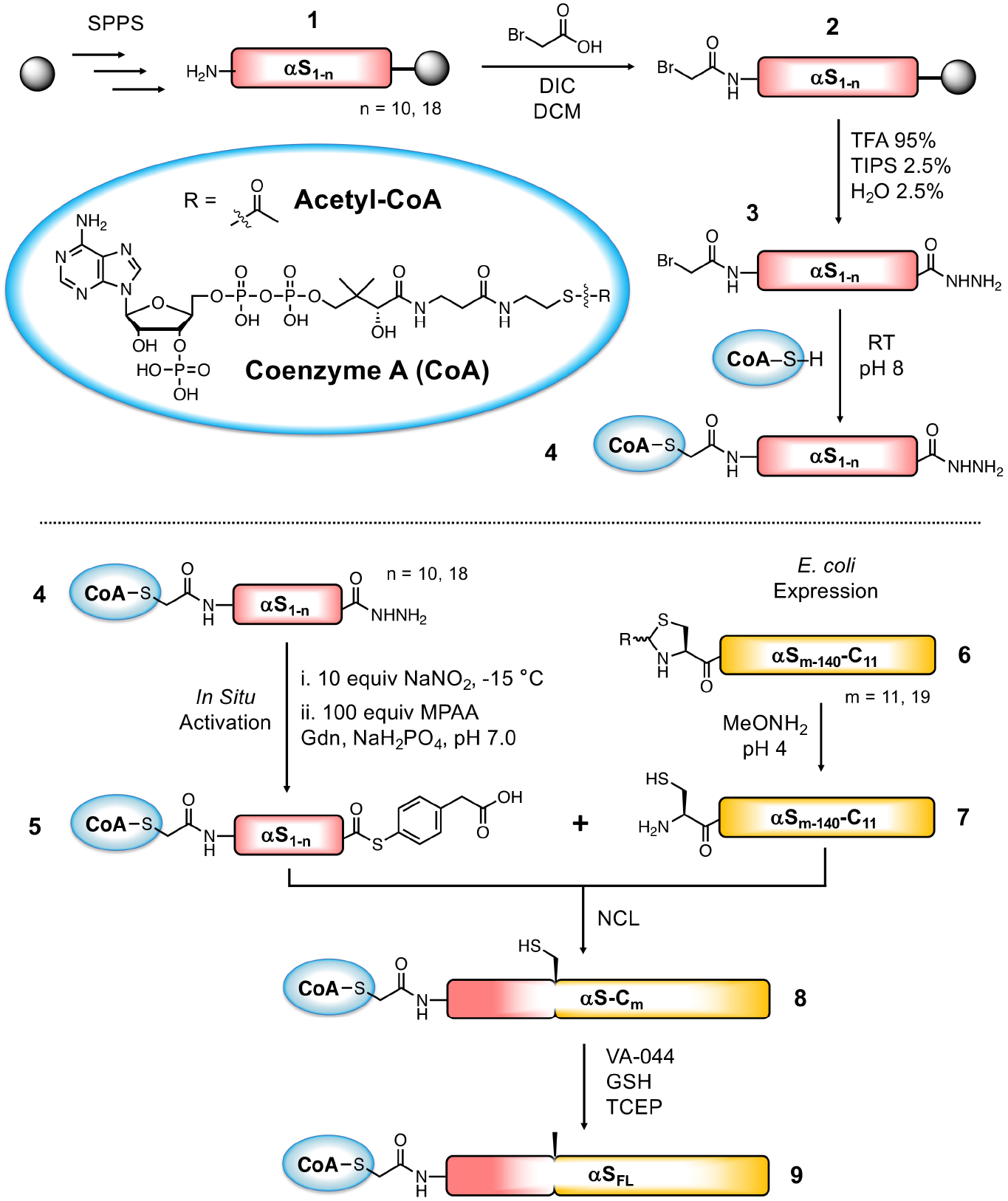
Semi-synthetic strategy for CoA-αS_FL_ inhibitor. Top: Synthesis of CoA-linked peptide acyl hydrazide fragment. Bottom: Semi-synthesis of CoA-αS_FL_ by native chemical ligation. Ligation at residues 10/11 (n = 10, m = 11) is described in detail in the main text, and compound numbers correspond to those fragments. Ligation at residues 18/19 (n = 18, m = 19) is described in the SI.

**Scheme 2.**
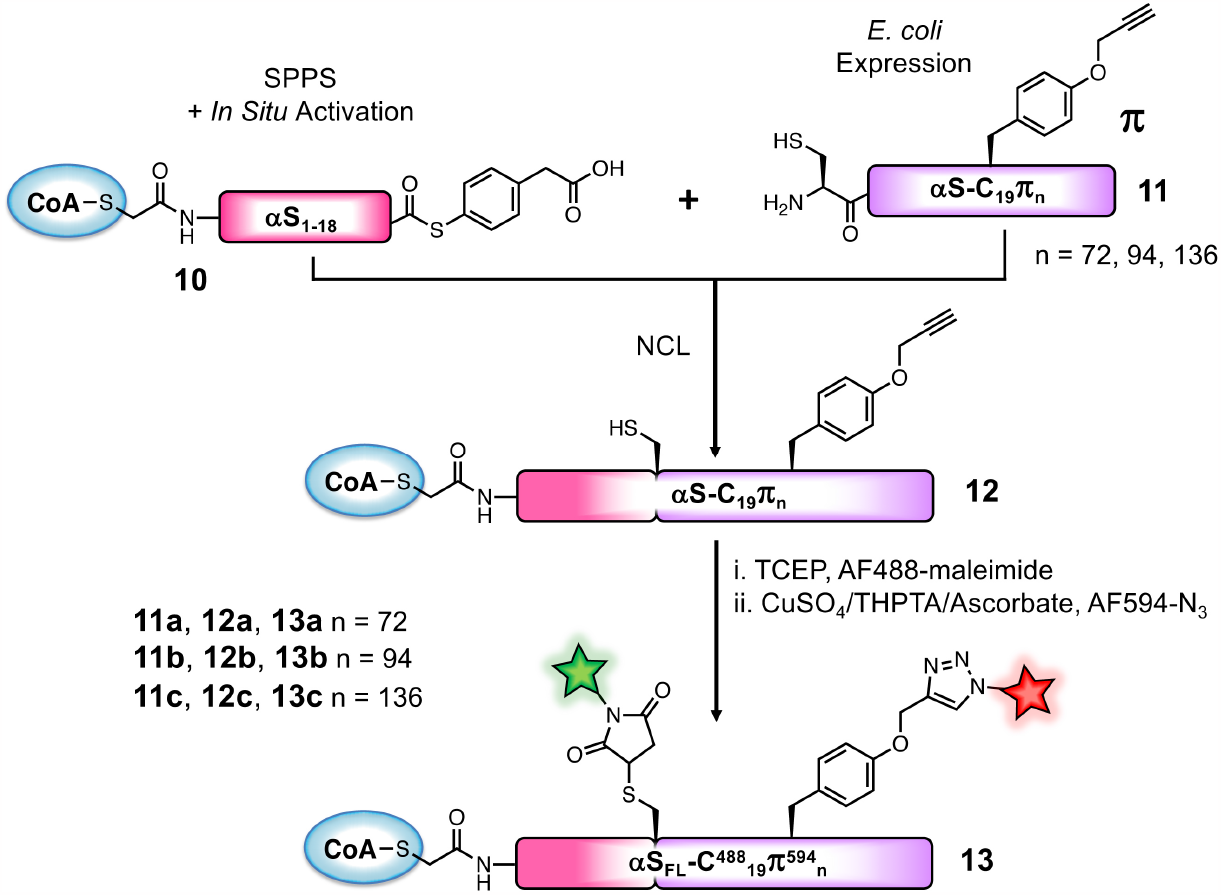
Semi-synthetic strategy for doubly fluorescently labeled inhibitor. MPAA thioester **10** is synthesized similarly to CoA-peptide **5** in Scheme 1. Protein fragments **11a, 11b**, and **11c** are obtained by unnatural amino acid incorporation during *E. coli* expression.

**Table 1.** smFRET efficiencies and histogram widths.

| Inhibitor or Complex | $ET_{\text{eff}}$ | Width |
| --- | --- | --- |
| $\alpha S(19-72)$ | $0.65 \pm 0.001$ | $0.15 \pm 0.003$ |
| $\alpha S(19-72) + \text{hNatB}$ | $0.64 \pm 0.023$ | $0.17 \pm 0.019$ |
| $\alpha S(19-94)$ | $0.48 \pm 0.001$ | $0.16 \pm 0.004$ |
| $\alpha S(19-94) + \text{hNatB}$ | $0.43 \pm 0.014$ | $0.24 \pm 0.010$ |
| $\alpha S(19-136)$ | $0.30 \pm 0.003$ | $0.21 \pm 0.002$ |
| $\alpha S(19-136) + \text{hNatB}$ | $0.25 \pm 0.005$ | $0.24 \pm 0.005$ |

Results shown as mean with standard deviation ( $n=3$ ). $\alpha S(19-n)$ refers to measurements made with CoA- $\alpha S$ -C<sup>488</sup><sub>19</sub> $\pi$ <sup>594</sup><sub>n</sub> (**13a**, **13b**, or **13c**).

