## Supporting Information for "Semi-synthetic CoA-α-Synuclein Constructs Trap N-terminal Acetyltransferase NatB for Binding Mechanism Studies"

**Contents:**

*List of figures and tables* ........ **S2**
*General information* ........ **S3**
*Synthesis of CoA-αS$_{1-10}$ (**4**) and CoA-αS$_{1-18}$ (**S2**) acyl hydrazides* ........ **S4**
*Expression and thiazolidine deprotection of αS$_{11-140}$-C$_{11}$ (7) and αS$_{19-140}$-C$_{19}$π$_n$* (**11a**, **11b**, **11c**) ........ **S5**
*Native chemical ligation* ........ **S6**
*Fluorescent Labeling of NCL products CoA-αS-C$_{19}$π$_n$ (**12a**, **12b**, **12c**)* ........ **S7**
*Purification of hNatB* ........ **S7**
*IC$_{50}$ determination of CoA- αS on hNatB activity* ........ **S8**
*Cryo-EM studies of hNatB and CoA-αS$_{FL}$ (**9**)* ........ **S8**
*Determining full binding of CoA-αS-C$_{19}$π$_n$ by fluorescence correlation spectroscopy* ........ **S9**
*Fraction of inhibitor bound to hNatB* ........ **S10**
*Characterizing CoA-αS and hNatB interactions by single molecule FRET* ........ **S11**
*Fluorescence polarization and anisotropy* ........ **S12**
*Simulations of CoA-αS bound to hNatB* ........ **S13**

**Figures:**

Figure S1. Synthetic peptide CoA-α$S_{1\text{-}10}$ with C-terminal hydrazide (**4**) .................................... **S15**
Figure S2. Recombinant Thz-α$S_{11\text{-}140}$-$C_{11}$ (**6**) .................................... **S16**
Figure S3. Deprotection of Thz-α$S_{11\text{-}140}$-$C_{11}$ (**6**) by methoxyamine .................................... **S16**
Figure S4. NCL of CoA-α$S_{1\text{-}10}$-$NHNH_2$ (**4**) and α$S_{11\text{-}140}$-$C_{11}$ (**7**) .................................... **S17**
Figure S5. Desulfurization of CoA-αS-$C_{11}$ (**8**) .................................... **S18**
Figure S6. Potency of hNatB inhibition by CoA-α$S_{1\text{-}10}$ and CoA-α$S_{FL}$ (**9**) .................................... **S18**
Figure S7. Additional images of hNatB + CoA-α$S_{FL}$ (**9**) Structure. .................................... **S19**
Figure S8. Recombinant α$S_{18\text{-}140}$-$C_{18}\pi_n$ (**S5a**, **S5b**, **S5c**) .................................... **S21**
Figure S9. Synthetic peptide CoA-α$S_{1\text{-}18}$ with C-terminal hydrazide (**S2**) .................................... **S22**
Figure S10. Synthetic peptide CoA-α$S_{1\text{-}18}$ with C-terminal alkyl thioester (**S3**) .................................... **S23**
Figure S11. Recombinant α$S_{19\text{-}140}$-$C_{19}$ (**S4**) .................................... **S23**
Figure S12. Recombinant α$S_{19\text{-}140}$-$C_{19}\pi_{72}$ (**11a**) .................................... **S24**
Figure S13. Recombinant α$S_{19\text{-}140}$-$C_{19}\pi_{94}$ (**11b**) .................................... **S25**
Figure S14. Recombinant α$S_{19\text{-}140}$-$C_{19}\pi_{136}$ (**11c**) .................................... **S26**
Figure S15. NCL of CoA-α$S_{1\text{-}18}$ (**S3**) and α$S_{19\text{-}140}$-$C_{19}$ (**S4**) .................................... **S27**
Figure S16. NCL of CoA-α$S_{1\text{-}18}$ (**10**) and α$S_{19\text{-}140}$-$C_{19}\pi_{72}$ (**11a**) .................................... **S28**
Figure S17. NCL of CoA-α$S_{1\text{-}18}$ (**10**) and α$S_{19\text{-}140}$-$C_{19}\pi_{94}$ (**11b**) .................................... **S28**
Figure S18. NCL of CoA-α$S_{1\text{-}18}$ (**10**) and α$S_{19\text{-}140}$-$C_{19}\pi_{136}$ (**11c**) .................................... **S29**
Figure S19. Fluorescent labeling of CoA-αS-$C_{19}\pi_{72}$ (**12a**) .................................... **S29**
Figure S20. Fluorescent labeling of CoA-αS-$C_{19}\pi_{94}$ (**12b**) .................................... **S30**
Figure S21. Fluorescent labeling of CoA-αS-$C_{19}\pi_{136}$ (**12c**) .................................... **S31**
Figure S22. Full binding of CoA-αS-$C^{488}{}_{19}\pi_n$ (**13a**, **13b**, **13c**) to NatB by FCS .................................... **S32**
Figure S23. Conformation of CoA-αS-$C^{488}{}_{19}\pi^{594}{}_{72}$ (**13a**) and complex with NatB by smFRET ... **S34**
Figure S24. Conformation of CoA-αS-$C^{488}{}_{19}\pi^{594}{}_{94}$ (**13b**) and complex with NatB by smFRET ... **S35**
Figure S25. Conformation of CoA-αS-$C^{488}{}_{19}\pi^{594}{}_{136}$ (**13c**) and complex with NatB by smFRET .. **S36**
Figure S26. Fluorescence polarization/anisotropy of CoA-αS-$C^{488}{}_{19}\pi_n$ (**13a**, **13b**, **13c**) ± NatB.... **S37**
Figure S27. Analysis of simulated αS/hNatB structures .................................... **S38**
Figure S28. Representative structures and energies from raw and refined αS/hNatB ensembles .. **S39**
Figure S29. αS/hNatB contact propensities .................................... **S40**
Figure S30. Simulated and experimental distance distributions for residues in smFRET .................................... **S41**

**Tables:**

Table S1. Summary statistics for hNatB/CoA-α$S_{FL}$ CryoEM Map .................................... **S20**
Table S2. FCS data .................................... **S32**
Table S3. smFRET data .................................... **S33**
Table S4. Fluorescence anisotropy data .................................... **S37**

*General Information*

Reagents for peptide synthesis, including 2-(1H-benzotriazol-1-yl)-1,1,3,3-tetramethyluronium hexafluorophosphate (HBTU), 7-azabenzotriazol-1-yloxy)tripyrrolidinophosphonium hexafluoro-phosphate (PyAOP), *N*,*N*-diisopropylethylamine (DIPEA), and Fmoc-amino acids, were purchased from EMD Millipore (Burlington, MA, USA). Coenzyme A sodium salt hydrate was purchased from Millipore Sigma (Burlington, MA, USA). Coenzyme A lithium salt was purchased from CoALA Biosciences (Elgin, TX, USA). DNA oligomers were purchased from Integrated DNA Technologies, Inc (Coralville, IA, USA). DNA extraction and Miniprep kits were purchased from Qiagen (Hilden, Germany). The starting plasmid pTXB1-αS-intein-$His_6$ containing αS with a C-terminal fusion to the *Mycobacterium xenopi* GyrA intein and C-terminal $His_6$ tag was constructed as described previously.[1] Buffers were made with MilliQ filtered (18 MΩ) water (Millipore; Billerica, MA, USA). *E. coli* BL21(DE3) cells were purchased from Stratagene (La Jolla, CA, USA). Reagents for native chemical ligation (NCL): $NaNO_2$, *tris*(2-carboxyethyl)phosphine (TCEP), and mercaptophenyl acetic acid (MPAA) were purchased from Sigma-Aldrich (St. Louis, MO, USA). Sodium mercaptoethanesulfonate (MESNa) was purchased from Tokyo Chemical Industry (Tokyo, Japan). Matrix-assisted laser desorption/ionization (MALDI) mass spectra were collected with a Ultraflex III MALDI-TOF/TOF mass spectrometer, a Microflex MALDI-TOF mass spectrometer, or a Rapiflex MALDI-TOF/TOF mass spectrometer (Bruker; Billerica, MA, USA). SDS-PAGE gels were imaged with a Typhoon FLA 7000 (GE Lifesciences; Princeton, NJ, USA). Peptides and protein fragments were purified on a PrepStar 218 module equipped with a ProStar 701 fraction collector and a ProStar 335 photodiode array detector (Varian; Walnut Creek, CA, USA) or a 1260 Infinity II preparative LC system (Agilent Technologies; Santa Clara, CA, USA). NCL reactions were monitored on a 1260 Infinity II LC system (Agilent Technologies; Santa Clara, CA, USA) using a Jupiter C4 column (Phenomenex; Torrance, CA, USA). Water + 0.1% trifluoroacetic acid (TFA) (solvent A) and acetonitrile + 0.1% TFA (solvent B) were used as the mobile phase in HPLC.

*Synthesis of CoA-αS$_{1-10}$ (**4**) and CoA-αS$_{1-18}$ (**S2**) acyl hydrazides*

Solid phase peptide synthesis (SPPS) of $H_2N$-αS$_{1-10}$ (**1**) was carried out using 2-chloro-trityl resin following standard procedures. The resin was derivatized by coupling Fmoc-hydrazine, and extra sites were capped with methanol. Couplings were carried out at a concentration of 0.2-0.3 M using 5 equiv Fmoc-protected amino acid, 5 equiv 7-azabenzotriazol-1-yloxy)tripyrrolidinophosphonium hexafluorophosphate (PyAOP), and 10 equiv DIPEA. The activated mixture was added to the deprotected peptidyl resin and stirred for 30 minutes. Washes with DMF and $CH_2Cl_2$ were performed between each coupling. Fmoc deprotection was done by adding 20% (v/v) piperidine in DMF and stirring for 20 min. $H_2N$-αS$_{1-18}$ (**S1**) was synthesized following the same protocol except using 4.9 equiv 2-(1H-benzotriazol-1-yl)-1,1,3,3-tetramethyluronium hexafluorophosphate (HBTU) and 4.9 equiv 1-hydroxybenzotriazole hydrate (HOBt). Both peptides were synthesized on a 100 μmol scale.

After removal of the final Fmoc group, bromoacetic acid was coupled as an active *O*-acylisourea using carbodiimide chemistry. 10 equiv bromoacetic acid and 9.3 equiv *N*,*N'*-diisopropylcarbodiimide were dissolved in DMF. The mixture was added to the peptidyl-resin and stirred at room temperature. Bromoacetylation proceeded to completion in 1 h.

Cleavage of bromoacetyl-αS$_{1-10}$-$NHNH_2$ (**3**) or bromoacetyl-αS$_{1-18}$-$NHNH_2$ (**S1'**) from the resin was performed using 30 μL of cleavage cocktail (95% TFA, 2.5% triisopropylsilane, 2.5% $H_2O$) per mg of resin and agitating for 1 h at room temperature. The cocktail containing the cleaved peptide was collected, and the peptide was precipitated using at least a 10-fold volume of diethyl ether. The precipitate was dissolved in $H_2O$/MeCN and lyophilized.

An aliquot of the lyophilized peptide **3** (6.8 mg, 5.3 μmol) was redissolved in triethylammonium bicarbonate (TEAB) buffer pH 8 to a final concentration of 10 mM. 2 equiv coenzyme A sodium salt hydrate was added, and the mixture was allowed to react for 4 h at room temperature and then overnight at 4 °C. Product formation was confirmed by MALDI-MS. The final product was purified by RP-HPLC over a C4 column. Characterization of CoA-αS$_{1-10}$-$NHNH_2$ (**4**) is shown in Figure S1.

The reaction of **S1'** with coenzyme A lithium salt at room temperature overnight also afforded the desired product. In this case, the resulting hydrazide was converted to the corresponding thioester for easy purification. Specifically, an aliquot of the lyophilized bromoacetyl-αS$_{1-18}$-$NHNH_2$ (**S1'**, 6.7 mg, 3.3 μmol) was redissolved in TEAB buffer pH 8 to a final concentration of 3 mM. Two equiv coenzyme A lithium salt was added, and the mixture was allowed to react at room temperature overnight. Product formation was confirmed by MALDI-MS. The alkylated product was roughly purified by RP-HPLC over a C4 column. Characterization of CoA-αS$_{1-18}$-$NHNH_2$ (**S2**) is shown in Figure S9. To a solution of the hydrazide **S2** (2.2 mg, 0.72 μmol) in low pH buffer (0.29 mL, 6 M guanidinium, 200 mM $NaH_2PO_4$, pH 3.0) was added 10 equiv $NaNO_2$ at -15 °C in an ice-salt bath. After 15 min reaction at -15 °C, 100 equiv MESNa was added to the mixture. The pH was adjusted to 7.0 and the reaction was let sit for 30 min at room temperature. The desired thioester was purified by RP-HPLC over a C4 semi-preparative column. Characterization of CoA-αS$_{1-18}$-$S(CH_2)_2SO_3H$ (**S3**) is shown in Figure S10.

*Expression and thiazolidine deprotection of αS$_{11-140}$-C$_{11}$ (7) and αS$_{19-140}$-C$_{19}\pi_n$ (**11a, 11b, 11c**)*

Deletion PCR was performed on plasmid containing full-length αS-Mxe-His$_6$ to generate the αS$_{11-140}$-C$_{11}$ (**7**) construct as a ligation partner for NCL. Protein fragment was expressed as previously described.[2] After Ni-NTA affinity column, the intein was cleaved from the αS fragment using 200 mM β-mercapto ethanol (βME) on a rotisserie overnight at room temperature. The resulting αS$_{11-140}$-C$_{11}$ with C-terminal carboxylate was dialyzed in buffer (20 mM Tris pH 8), purified by a second Ni-NTA column to remove the cleaved intein, and further purified by fast protein liquid chromatography (FPLC) over a 5 mL Hi-Trap Q column. Expression of αS C-terminal fragments in *E. coli* results in its production with the initiator methionine, which is cleaved *in vivo* by endogenous methionine amino peptidase (MAP).[3] Endogenous aldehydes react with the exposed N-terminal cysteine to form thiazolidine adducts (Thz).[4] Therefore the Thz[5] was removed using 100 mM methoxyamine at pH 4 to regenerate the N-terminal cysteine for later use in ligation. Deprotected fragment was purified by RP-HPLC using a C4 column.  Expression and deprotection data are shown in Figures S2 and S3. Plasmid containing the αS$_{19-140}$-C$_{19}$ construct was generated by deletion PCR using the primers shown below. Expression and purification were performed as described above. Characterization of **S4** is shown in Figures S11.

Unnatural amino acid incorporation via amber codon suppression was used to produce αS$_{18-140}$-C$_{18}\pi_n$ (n = 72, 94, or 136; **S3a**, **S3b**, **S3c**, respectively) and αS$_{19-140}$-C$_{19}\pi_n$ (n = 72, 94, or 136; **11a**, **11b**, **11c**, respectively). Plasmid containing the desired αS construct was generated by deletion PCR and transformed into *E. coli* pDULE-pXF cells with pre-transformed plasmids encoding propargyl-tyrosine (Ppy or π) synthetase and tRNA. Expression was carried out as above, except cells were grown in M9 minimal media, and π (220 mg/L) was added to the culture at OD ~0.8 with a 10-min incubation prior to inducing expression with IPTG. Deprotection of Thz from $C_{19}$ and purification of $\alpha S_{19\text{-}140}$-$C_{19}\pi_n$ (n = 72, 94, or 136; **11a**, **11b**, **11c**, respectively) were carried out as for $\alpha S_{11\text{-}140}$-$C_{11}$ (**7**). Characterizations of **11a**, **11b**, **11c** are shown in Figures S10, S11, and S12, respectively.

Primer sequences:

| | | |
|---|---|---|
| $\alpha S_{11\text{-}140}$ | Forward | 5'-TGCAAGGAGGGAGTTGT-3' |
| | Reverse | 5'-CATATGTATATCTCCTTCTTAAAGTTAAAC-3' |
| $\alpha S_{18\text{-}140}$ | Forward | 5'-TGTGCTGAGAAAACCAAACAG-3' |
| | Reverse | 5'-CATATGTATATCTCCTTCTTAAAGTTAAAC-3' |
| $\alpha S_{19\text{-}140}$ | Forward | 5'-TGTGAGAAAACCAAACAGGG-3' |
| | Reverse | 5'-CATATGTATATCTCCTTCTTAAAGTTAAAC-3' |

*Native chemical ligation*

CoA-$\alpha S_{1\text{-}10}$ acyl hydrazide (**4**) was dissolved in low pH NCL buffer (6 M guanidinium, 200 mM $NaH_2PO_4$, pH 3) for a final concentration of 2 mM and chilled to -15 °C in an ice-salt bath. Hydrazide to azide conversion was achieved by adding 10 equiv $NaNO_2$ and agitating by magnetic stirring for 15 min at -15 °C. 40 equiv MPAA, pre-dissolved in NCL buffer pH 7.0, was added to the mixture, followed by 2 equiv partner fragment $\alpha S_{11\text{-}140}$-$C_{11}$. The reaction was warmed to room temperature, and the pH was adjusted to 7.0. TCEP was added to 40 mM final concentration, and the reaction was incubated at 37 °C with agitation at 500 rpm. Product formation was monitored by MALDI-MS and analytical HPLC. To convert Cys11 used in ligation to the native alanine, the purified NCL product was re-dissolved in pH 7 NCL buffer and incubated with 20 mM radical initiator VA-044, 100 mM glutathione (GSH), and 250 mM TCEP in an argon-purged tube at 37 °C overnight. The desulfurized full-length product was purified by RP-HPLC over a C4 semi-preparative column. NCL and desulfurization reactions are shown in Figures S4 and S5, respectively.

Alternatively, the same product CoA-$\alpha S_{FL}$ (**9**) was synthesized through the NCL of CoA-$\alpha S_{1\text{-}18}$-MES (**S3**, 0.22 µmol) and 1.8 equiv $\alpha S_{19\text{-}140}$-$C_{19}$ (**S4**, 0.40 µmol). These fragments were dissolved in NCL buffer (0.22 mL, 6 M guanidinium, 200 mM $Na_2HPO_4$, 100 mM MPAA, 50 mM TCEP, pH 7) and the reaction was incubated at 37 °C. Product formation was monitored by MALDI-MS and analytical HPLC. After completion of the NCL, the pH was adjusted to 2 and MPAA was extracted with $Et_2O$ thoroughly. The residual $Et_2O$ was removed by argon stream and the pH was adjusted back to 7. To the resulting solution was added desulfurization buffer (0.75 mL, 6 M guanidinium, 200 mM $Na_2HPO_4$, 500 mM TCEP, 200 mM MESNa, pH 7) and 100 equiv VA-044. After incubation at 37 °C overnight, the desulfurized product was purified by RP-HPLC over a C4 semi-preparative column. NCL and desulfurization reactions are shown in Figures S15.

NCLs for CoA-αS-$C_{19}\pi_{72}$ (**12a**), CoA-αS-$C_{19}\pi_{94}$ (**12b**), and CoA-αS-$C_{19}\pi_{136}$ (**12c**) were carried out as above using CoA-α$S_{1\text{-}18}$ (**S2**) and 1.2 equiv C-terminal partner fragment. Instead of desulfurization, the Cys19 of the ligation product was labeled with Alexa Fluor 488 (AF488) maleimide. Each NCL was done on a ~300 nanomole scale. NCL data are shown in Figures S13-S15.

*Fluorescent Labeling of NCL products CoA-αS-$C_{19}\pi_n$ (**12a, 12b, 12c**)*

NCL product CoA-αS-$C_{19}\pi_{72}$ (**12a**), CoA-αS-$C_{19}\pi_{94}$ (**12b**), or CoA-αS-$C_{19}\pi_{136}$ (**12c**) was redissolved in buffer (2 M guadidinium, 67 mM $NaH_2PO_4$, pH 7.0) and incubated with 2 equiv tris(2-carboxyethyl)phosphine (TCEP) at 37 °C for ~10 minutes. The fragments were labeled by adding 2 equiv AF488-maleimide pre-dissolved in DMSO. The reaction tube was wrapped in aluminum foil and incubated at 37 °C for 3 h or until completion of labeling, monitored by MALDI-MS. Labeled proteins were purified by reverse-phase high-performance liquid chromatography (RP-HPLC) using a C4 column. To carry out the second labeling with Alexa Fluor 594 (AF594), the purified singly labeled product CoA-αS-$C^{488}{}_{19}\pi_{72}$, CoA-αS-$C^{488}{}_{19}\pi_{94}$, or CoA-αS-$C^{488}{}_{19}\pi_{136}$ was re-dissolved in buffer (0.2 M guanidinium, 0.2 M $NaH_2PO_4$, pH 7.0). Catalytic mixture consisting of 2 equiv $CuSO_4$, 10 equiv THPTA, and 20 equiv sodium ascorbate was let sit for 10 min and added to the protein along with 2 equiv fluorophore. Labeled proteins were supplemented with TCEP before purification by RP-HPLC. Labeling reaction data are shown in Figures S16-S18.

*Purification of hNatB*

hNatB was expressed in sf9 cells and purified as described previously.[6] Cells were harvested 48 h post infection and lysed by sonication in lysis buffer containing 25 mM Tris, pH 8.0, 300 mM NaCl, 10 mM Imidazole, 10 mM βME, 1 mM phenylmethylsulfonyl fluoride (PMSF) and one ethylenediamine tetraacetic acid- (EDTA-) free protease inhibitor tablet. Clarified lysate was passed onto the Ni-NTA resin (Thermo Scientific), washed extensively with lysis buffer, and eluted with lysis buffer containing 300 mM imidazole. Protein was dialyzed into buffer containing 25 mM HEPES, pH 7.5, 50 mM NaCl, 10 mM βME for SP column ion-exchange and eluted with a salt gradient (50–750 mM NaCl). Peak fraction was further concentrated for a S200 gel-filtration run in a buffer with 25 mM HEPES, pH 7.5, 200 mM NaCl, and 1 mM TCEP. Purified protein samples were snap-frozen in liquid nitrogen and stored at -80°C for further use.

*$IC_{50}$ inhibition assay of CoA-α$S_{FL}$ on hNatB activity*

hNatB acetyltransferase assays were carried out at room temperature in a reaction buffer containing 75 mM HEPES, pH 7.5, 120 mM NaCl, 1 mM DTT as described previously.[6] 100 nM hNatB was mixed with 500 μM 'MDVF' peptide ($NH_2$-MDVFMKGRWGRPVGRRRRP-COOH, GeneScript) and 300 μM $^{14}C$ labeled acetyl-CoA ($^{14}C$-labeled, 4 mCi mmol$^{-1}$; PerkinElmer Life Sciences), and CoA-αS concentrations were varied (0 μM, and 0.23 μM to 13.44 μM) for a 10-minute reaction. Reactions were quenched by adding the solution onto P81 paper discs (St. Vincent's Institute Medical Research). P81 paper discs were washed extensively to remove unreacted $^{14}C$ labeled acetyl-CoA and then subjected to radioactive counts (Packard Tri-Carb 1500 liquid scintillation analyzer). Data points (Fig. S6) were normalized to reaction with no CoA-αS added and fitted to a sigmoidal dose-response curve with GraphPad Prism (version 5.01). Errors represent s.d. (n = 3).

*Cryo-EM studies of hNatB and CoA-α$S_{FL}$ (**9**)*

To prepare the single particle analysis cryo-EM sample, purified hNatB was mixed with 2-fold excess of CoA-α$S_{FL}$ (**9**) and concentrated to 4 mg/mL. 20μL of this sample supplemented with 0.05% of Tween-20 was applied to glow-discharged Quantinfoil R1.2/1.3 holey carbon support grids, blotted and plunged into liquid ethane using an FEI Vitrobot Mark IV. Data collection was performed with a Titan Krios equipped with a K3 Summit direct detector (Gatan) at a magnification of 105,000 with defocus values from -1.0 to -3.0 μm and a total dose of 43.9 e-/Å$^2$, resulting in 40 frames per stack. Image stacks were automatically collected with Latitude software (Gatan, Inc). Image stacks were corrected for beam induced motion using MotionCor2[7] and binned two fold resulting in a pixel size of 0.86Å. CTFFIND4[8] was utilized for CTF estimation. A pretrained Topaz model,[9] ResNet16, was utilized for particle picking and resulted in a total of 741,341 particles. Particles were extracted and low pass filtered to 12Å for 2D classification. Multiple rounds of homogeneous and heterogeneous refinement were utilized to remove bad particles. The final map was post-processed using DeepEMhancer[10] for visualization purposes. The resulting map has an overall resolution of 3.39 Å. Map quality was assessed using both 3DFSC[11] and CryoEF.[12] The model was built from a previously reported cryoEM structure of hNatB with $\alpha S_{1-10}$ (PDB: 6VP9) in COOT[13] and real space refined in Phenix.[14] The entire CMC ligand was not resolved in the structure, so we therefore modeled a truncated form of the ligand into the map. Additionally, there was no reliable cryo-EM density beyond $\alpha$S $Met_5$. The cryo-EM workflow is shown in Figure S7. The model and map were deposited to PDB (8G0L) and EMDB (29657).

*Determining full binding of CoA-αS-$C_{19}\pi_n$ by fluorescence correlation spectroscopy (FCS)*

FCS measurements were carried out using a lab-built setup as described in our previous works.[15-16] The instrument is based on an Olympus IX71 microscope with a continuous emission 488 nm DPSS 50 mW laser (Spectra-Physics, Santa Clara, CA). All measurements were made at 25 °C. The laser power entering the microscope was adjusted to 4.5 μW. Fluorescence emission collected through the objective was separated from the excitation signal through a Z488rdc long pass dichroic filter and an HQ600/200m bandpass filter (Chroma, Bellows Falls, VT). Emission signal was focused onto the aperture of a 50 μm optical fiber. Signal was detected by an avalanche photodiode (Perkin Elmer, Waltham, MA) directly coupled to the fiber. A digital autocorrelator (Flex03Q-12, correlator.com, Bridgewater, NJ) was used to collect 10 autocorrelation curves of 10 seconds for each measurement of CoA-αS free in buffer or as a complex with NatB. Fitting was done using lab-written code in MATLAB (The MathWorks, Natick, MA).

Eight-well chambered Nunc coverglasses (Thermo Fisher Scientific, Waltham, MA) were prepared by plasma cleaning followed by incubation overnight with polylysine-conjugated polyethylene glycol (PEG-PLL), prepared using a modified Pierce PEGylation protocol (Pierce, Rockford, IL). PEG-PLL coated chambers were rinsed with and stored in Milli-Q water until use.

To determine the diffusion time of the inhibitor, CoA-αS singly labeled with AF488 was measured in buffer (75 mM HEPES, 120 mM NaCl, 1 mM DTT, pH 7.4). The average of 10 autocorrelation curves was fit to a 1-component autocorrelation function:

$$G(\tau) = \frac{1}{N}\left(\frac{1}{1+\frac{\tau}{\tau_1}} * \left(\frac{1}{1+\frac{s^2\tau}{\tau_1}}\right)^{1/2}\right)$$

where $G(\tau)$ is the autocorrelation function, N is the number of molecules in the focal volume, $\tau_1$ is the diffusion time of CoA-αS, and s is the radial:axial ratio of the focal volume dimensions. The counts per molecule (CPM) for each sample was calculated by dividing the average intensity (Hz) of the measured signal by the number of molecules N. The normalized CPM of each CoA-αS was calculated by dividing by the CPM of freely diffusing fluorescent standard AF488. FCS data are shown in Table S1.

*Fraction of inhibitor bound to hNatB*

Enzyme-inhibitor complexes were formed by incubating singly labeled CoA-αS with various molar equivalents of NatB (1, 10, 40, or 100 equiv). The diffusion time of each complex was determined by fitting the average of 10 autocorrelation curves using the single component equation above. Saturation of inhibitor binding was confirmed as the diffusion time reached a plateau with increasing equiv NatB. The fraction of inhibitor bound to NatB was calculated by fitting each average autocorrelation curve to a 2-component equation:

$$G(\tau) = \frac{1}{N}\left(A * \frac{1}{1+\frac{\tau}{\tau_1}} * \left(\frac{1}{1+\frac{s^2\tau}{\tau_1}}\right)^{1/2} + (1-A) * \frac{1}{1+\frac{\tau}{\tau_2}} * \left(\frac{1}{1+\frac{s^2\tau}{\tau_2}}\right)^{1/2}\right)$$

where $G(\tau)$ is the autocorrelation function, N is the number of molecules in the focal volume, $\tau_1$ is the characteristic diffusion time of CoA-αS, $\tau_2$ is the characteristic diffusion time of the CoA-αS /NatB complex, s is the radial:axial ratio of the focal volume dimensions, and A is the fraction of CoA-αS free. $\tau_1$ was fixed to the diffusion time of CoA-αS free in solution, determined as described, and $\tau_2$ was fixed to the maximum diffusion of the complexes, determined from fitting to the single component equation. In each complex with different equivalents of NatB, the fraction of CoA-αS bound was obtained from fitting the autocorrelation curves collected using the complex. The resulting binding curve was fit to the following equation:

$$1 - A = \frac{B_{max}x}{K_{d,app} + x}$$

where $1 - A$ is the fraction of CoA-αS bound, x is the equivalents of NatB, $B_{max}$ is the maximum fraction of CoA-αS bound, and $K_{d,app}$ is the apparent dissociation constant. Averages of diffusion times etc. and standard deviations were calculated from at least 3 independent measurements. Binding data are shown in Figure S19.

*Characterizing CoA-αS and hNatB interactions by single molecule FRET*

All smFRET measurements were made on a MicroTime 200 inverse time- resolved confocal microscope (PicoQuant, Berlin, Germany). Eight-chambered Nunc coverslips (Thermo Fisher Scientific, Waltham, MA) were plasma cleaned and coated with PEG-PLL overnight, followed by rinsing with water prior to use. For each smFRET measurement, buffer (25 mM HEPES, 120 mM NaCl, pH 7.5) was added to a chamber to check for background signal, followed by measurement of sample in the same well. For measurements of inhibitor alone, 30 pM CoA-αS-$C^{488}{}_{19}\pi_n$ (**13a**, **13b**, **13c**) was used. For measurement of inhibitor in complex with enzyme, the complex was pre-formed by incubating CoA-αS-$C^{488}{}_{19}\pi_n$ (**13a**, **13b**, **13c**) with 100 equiv NatB, followed by dilution into the well. 485 nm and 561 nm lasers pulsed at 40 MHz were adjusted to 30 μW before entering the microscope. Fluorescence was collected through the objective and passed through a 150 μm pinhole. Excitation and emission were discriminated by passing the photons through a HQ585LP dichroic in combination with ET525/50M and HQ600LP filters. Signal was detected by photodiodes. Photon traces were collected in 1-ms time bins for an hour. A threshold of 30 counts/ms total in the donor and acceptor channels was used to discriminate events from noise. For each event, the energy transfer efficiency between donor and acceptor fluorophore ($ET_{eff}$) was calculated in the SymPhoTime 64 software using the following equation:

$$ET_{eff} = \frac{I_a - \beta I_d}{(I_a - \beta I_d) + \gamma(I_d + \beta I_d)}$$

where $I_a$ and $I_d$ are, respectively, the intensity of fluorescence detected in the acceptor and donor channels. β is the leakage of the fluorescence from the donor fluorophore into the acceptor channel, measured before each set of experiments. γ is the differences in detection efficiency and quantum yield between acceptor and donor fluorophores and is measured every few months. The resulting histograms were fit using Origin (OriginLab Corp, Northampton, MA) to Gaussian distributions:

$$y = \frac{A}{w\sqrt{\frac{\pi}{2}}} e^{-2\left(\left(\frac{(x-x_c)}{w}\right)^2\right)}$$

where w is the width, A is the area, and $x_c$ is the center of the distribution. FRET data are shown in Table S2 and Figures S20-S22.

*Fluorescence polarization and anisotropy*

For fluorescence polarization and anisotropy measurements, 50 µL of each sample was prepared in triplicate and pipetted into a Grenier black, clear flat bottom, half-area 96-well plate. For measurements of inhibitor alone, each AF594-labeled CoA-αS was diluted to 10 nM in buffer (25 mM HEPES, 120 mM NaCl, pH 7.5). For measurements of the inhibitor in complex with NatB, the complex was pre-formed by incubating CoA-αS with 100 equiv NatB, followed by dilution to 10 nM in buffer. Measurements were taken using a Tecan (Mannedorf, Switzerland) Spark plate reader. Excitation and emission monochromators were respectively set to 561 ± 20 nm and 624 ± 20 nm for AF594. For each sample, fluorescence polarization ratio was calculated using the following equation:

$$p = \frac{I_{\parallel} - GI_{\perp}}{I_{\parallel} + GI_{\perp}}$$

where p is the polarization ratio, $I_{\parallel}$ is the parallel intensity, $I_{\perp}$ is the perpendicular intensity, and G is the grating factor, an instrument correction factor calibrated using 10 nM AF594 free dye.

Similarly, anisotropy was calculated using the following equation:

$$r = \frac{I_{\parallel} - GI_{\perp}}{I_{\parallel} + 2GI_{\perp}}$$

where r is the anisotropy, $I_{\parallel}$ is the parallel intensity, $I_{\perp}$ is the perpendicular intensity, and G is the grating factor. Polarization and anisotropy data are shown in Table S3 and Figure S23.

*Simulations of CoA-αS bound to hNatB*

The hNatB with CoA-αS inhibitor bound cryo-EM structure was modified before use in FastFloppyTail (FFT).[17] To begin reformatting the structure for FFT input, the remaining unmodeled 135 residues of αS were built using PyMOL. The resulting structure was renumbered and the chain identifiers altered so that αS and hNatB were chains A and B/C, respectively, as required for FFT input. Finally, the sidechains of αS-hNatB were relaxed using the FastRelax. The hNatB backbone torsion angles were left unoptimized as this changed the structure significantly. Seqeunce-based secondary structure and disorder predictions were obtained from the FFT github (https://github.com/jferrie3/AbInitioVO-and-FastFloppyTail) and used to generate fragments from the Rosetta fragment picker utilizing the quota protocol as detailed by Fierre *et al.*[17]

Small adjustments to FFT were made to simulate αS in the presence of NatB. The option --Diso_Seg_Cutoff was added to specify the needed length of consecutive disordered residues to be considered a disordered region. A value of 4 was provided to allow the first 5 residues to remain constant. Additionally, the --Fold_Tree option was provided to allow propagation of the cryo-EM structure to the FFT simulated αS. Rotamer sampling of NatB was computationally expensive and induced artificial structural clashes since backbone motions were restricted. Therefore, the initial rotamers were saved from the starting structure to the initial 10,000 structures which afforded a simulation time of 10 minutes per structure. Utilizing these modifications, the final command used for structure generation was:

```
run FastFloppyTail.py -ftnstruct 10000 -t_frag asyn_quota_frags.200.3mers fold
5,1,-1.5,140,-1.5,277,1.277,1223,-1.277,141,-1 -diso
RAPX_REWEIGHTED_rapx.diso.txt -d_seg_cut 4 -inpdb Final_NatB.pdb
```

After selecting the 1000 lowest scoring decoys from the initial 10,000, the average final energy score using score function ref2015 was 2462 ± 16.9 Rosetta energy units (REU). After relaxing backbone and sidechain rotamers for αS and sidechain rotamers for NatB, the final score was -1579 ± 17.9 REU (ΔREU = -4041 REU). The energy distribution for the 1000 lowest energy structures is shown in Figure S24 along with the contact map for this ensemble, which we refer to as the raw simulated ensemble or raw ensemble. We note that a traditional "energy funnel" is not observed for this ensemble, which we believe to be a consequence of simulating disordered proteins due to many states occupying similar energy levels. The contact maps and energy distributions for free αS can be found in Ferrie et al.[18]

The distance distributions for the residue pairs used in smFRET experiments (19-72, 19-94, and 19-136) were calculated for the ensemble of 1000 lowest energy structures of free αS from Ferrie *et al.*[18] and the 1000 lowest energy structures of the αS/hNatB complex generated here. These are shown in Figure S25 along with experimental distance distributions for the αS/hNatB complexes derived from the smFRET distributions using a Gaussian chain model.[19-20]

The refined ensemble was generated using an additional score term to reweight scores for structures from the raw ensemble. Restraints on distances were modeled using the following equation:

$$s = k * (x - x_0)^2$$

Where x is the distance between $C_\alpha$ atoms in the pose, and $x_0$ is the distance derived from smFRET measurements. The variable k is a constant which was set to the standard deviation of the smFRET sample divided by 12. This score term was added to the total original score from the pose for the three pairs of residues for which smFRET data were available, 19-72, 19-94, 19-136. Those poses below zero REU were chosen as members of the refined ensemble.

A difference heat map showing the changes in the average inter-residue distances between the αS and refined αS/hNatB ensembles is shown in Figure 5 in the main tex. The corresponding map for the raw αS/hNatB ensemble is shown in Figure S27. The Rg of the refined αS/hNatB ensemble is compared to the Rg of the ensemble of free αS in Figure 5 in the main text. The Rg distributions for the raw and refined αS/hNatB ensembles are compared in Figure S27, as are the αS contact maps for each ensemble. Figure S28 compares the 10 lowest energy structures in each of the two αS/hNatB ensembles, and shows "energy funnel" plots with the distribution of energies of the structures in the two ensembles plotted against their root mean squared deviation (RMSD) from the lowest energy structure. Figure S29 compares the distance distributions from the smFRET experiments to simulations of αS alone and αS/hNatB in the raw and refined ensembles.

To determine whether any specific region of hNAA20 or hNAA25 had a significant level of contact with αS, a python script was used to cull any structures that placed αS residues within 10 Å of hNatB in at least 5% of structures in the ensemble. Figure S30 shows the hNatB structure with the region of highest contact, NAA25 residues 13-20, highlighted and plots showing the frequency of contacts for the most highly interaction portions of hNatB. This region is also highlighted in Figure 5 in the main text.

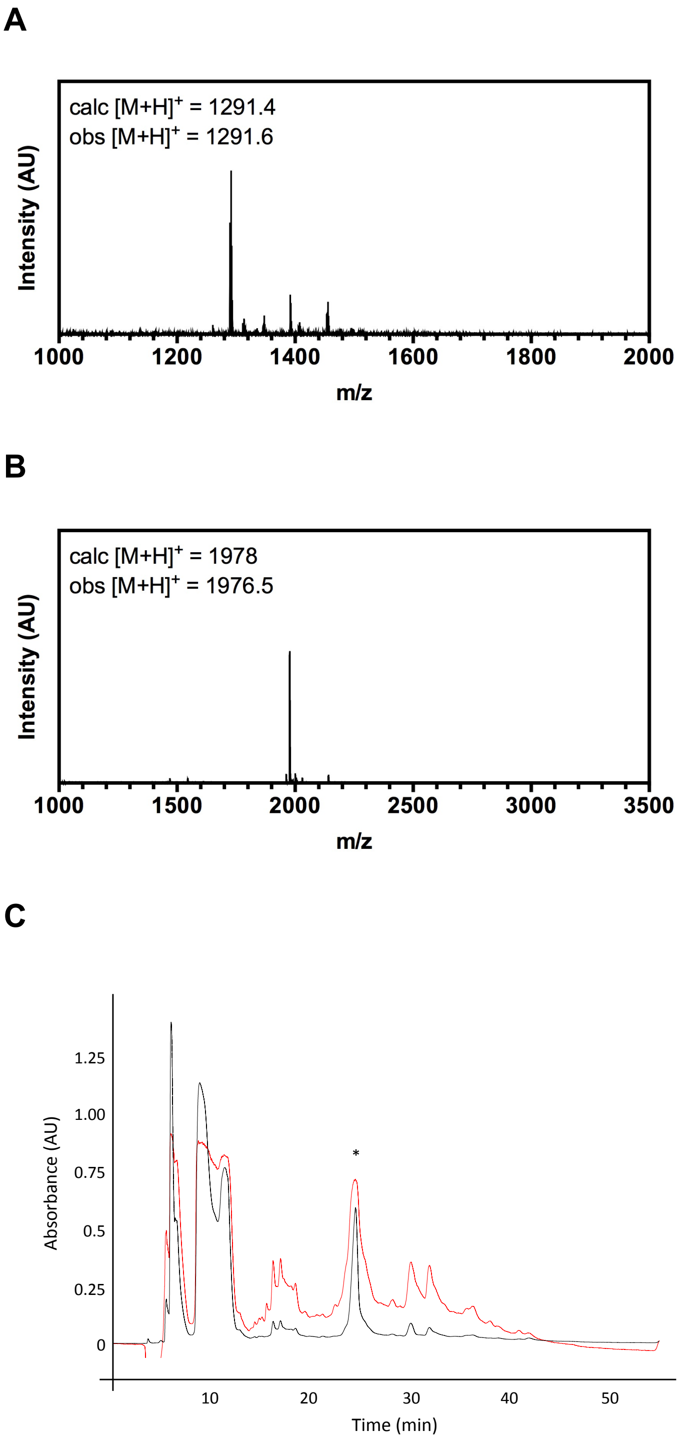

**Figure S1. Synthetic peptide CoA-αS$_{1-10}$ with C-terminal hydrazide (4). (A)** MALDI MS of Br-Ac-αS$_{1-10}$ **(B)** MALDI MS of CoA-αS$_{1-10}$ **(C)** Preparative HPLC (20-40% B over 30 min), red = 215 nm, black = 277 nm, * = product CoA-αS$_{1-10}$

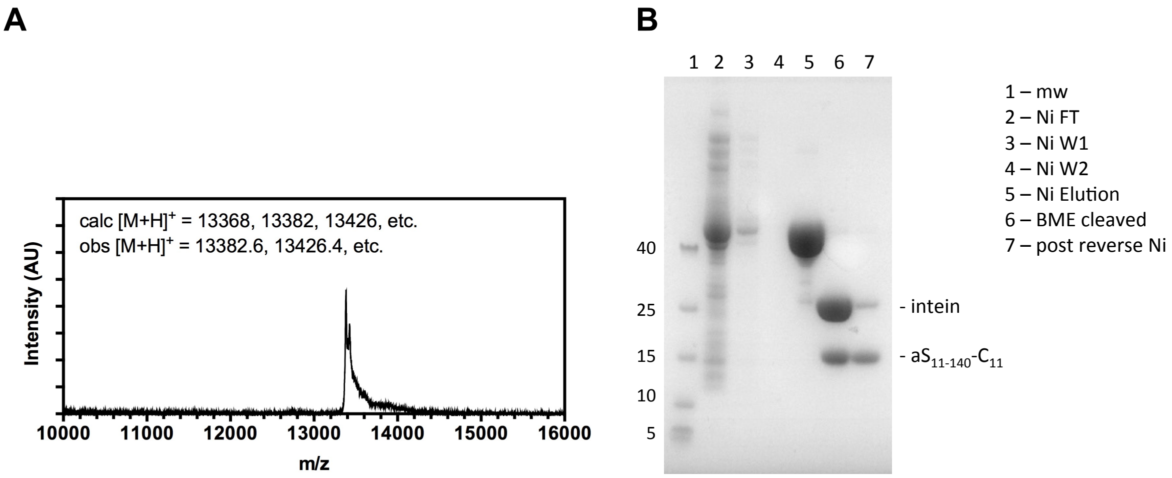

**Figure S2. Recombinant Thz-αS$_{11\text{-}140}$-C$_{11}$ (6). (A)** MALDI MS of purified Thz-αS$_{11\text{-}140}$-C$_{11}$ starting material. Observed masses correspond to various aldehyde- and ketone- derived thiazolidines. **(B)** SDS-PAGE (Coomassie stain)

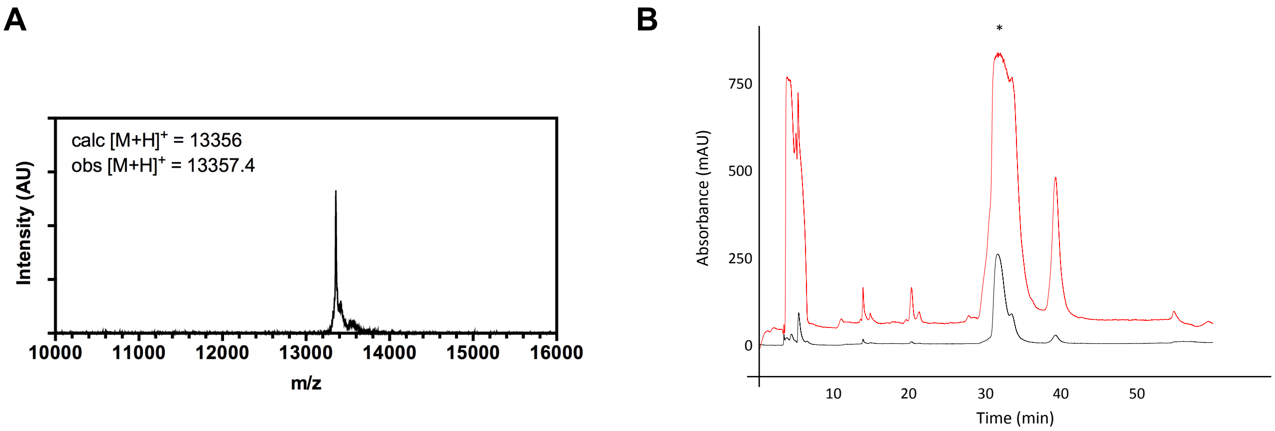

**Figure S3. Deprotection of Thz-αS$_{11\text{-}140}$-C$_{11}$ (6) by methoxyamine. (A)** MALDI MS of purified deprotection reaction. Observed mass corresponds to species with free N-terminus. **(B)** HPLC (25-50% B over 30 min), red = 215 nm, black = 277 nm, * = product

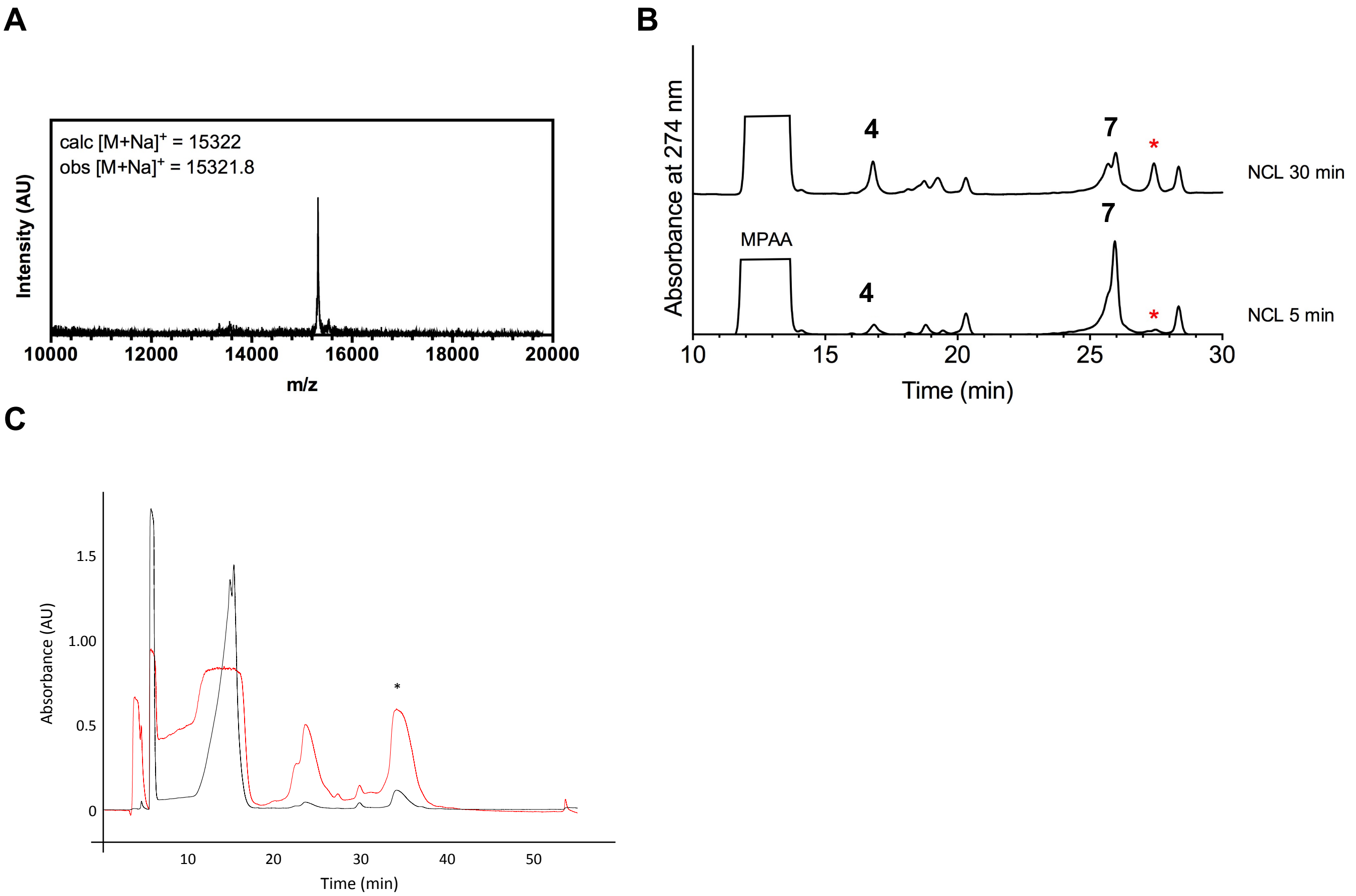

**Figure S4. NCL of CoA-αS$_{1-10}$-NHNH$_2$ (4) and αS$_{11-140}$-C$_{11}$ (7).** **(A)** MALDI MS of NCL product **(B)** Analytical HPLC (10-50% B over 30 min) of NCL reaction; CoA-αS$_{1-10}$-NHNH$_2$ (**4**), αS$_{11-140}$-C$_{11}$ (**7**), and product (*) (Retention time 27.4 min). **(C)** Preparative HPLC purification of NCL product (33-43% B over 30 min), red = 215 nm, black = 277 nm, * = product

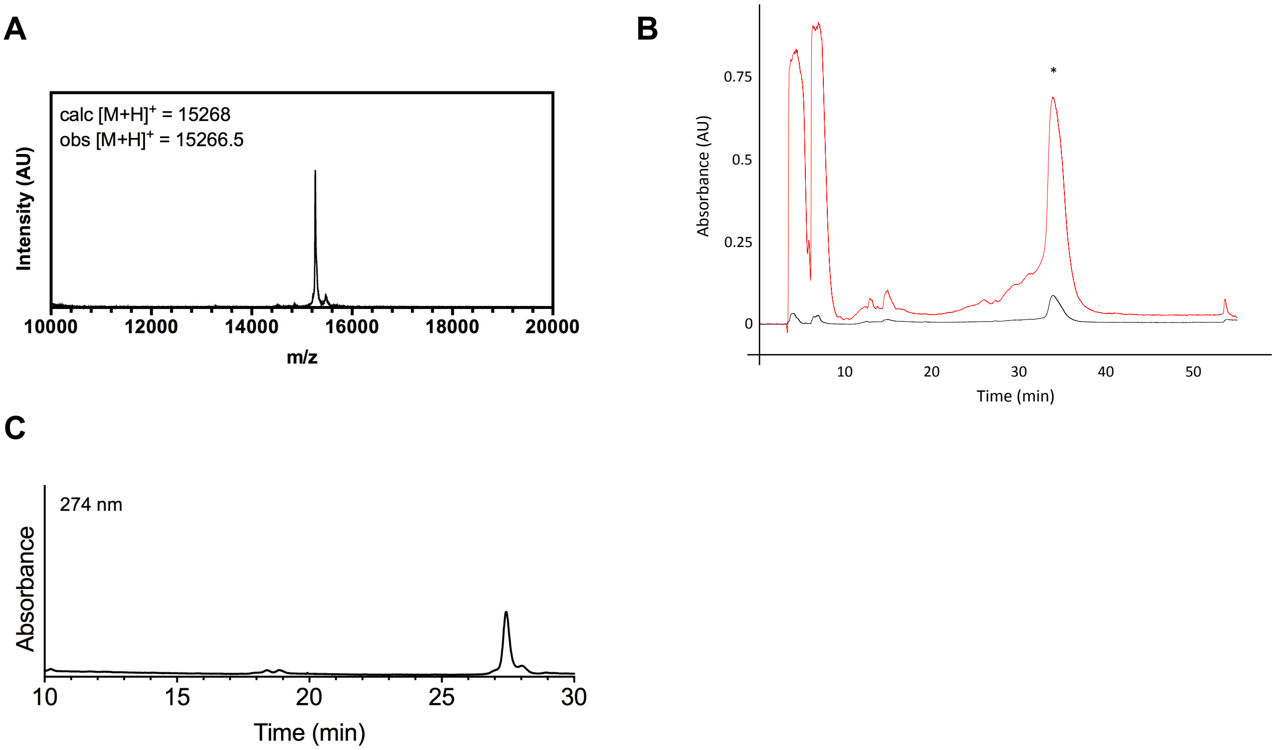

**Figure S5. Desulfurization of CoA-αS-$C_{11}$ (8). (A)** MALDI MS of desulfurized product CoA-αS **(B)** Preparative HPLC purification of desulfurized product (33-43% B over 30 min), red = 215 nm, black = 277 nm, * = product **(C)** Analytical HPLC (10-50% B over 30 min) of purified CoA-αS.

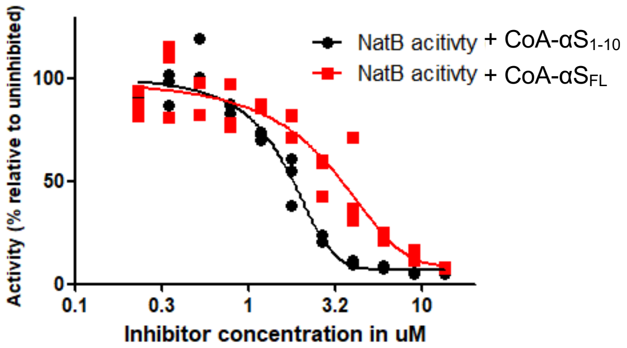

**Figure S6. Potency of hNatB inhibition by CoA-α$S_{1-10}$ and CoA-α$S_{FL}$ (9).** Dose-response curves representing the titration of CoA-α$S_{FL}$ (red) into hNatB acetyltransferase reactions, compared with that of CoA-α$S_{1-10}$ (black), determined previously.[6]

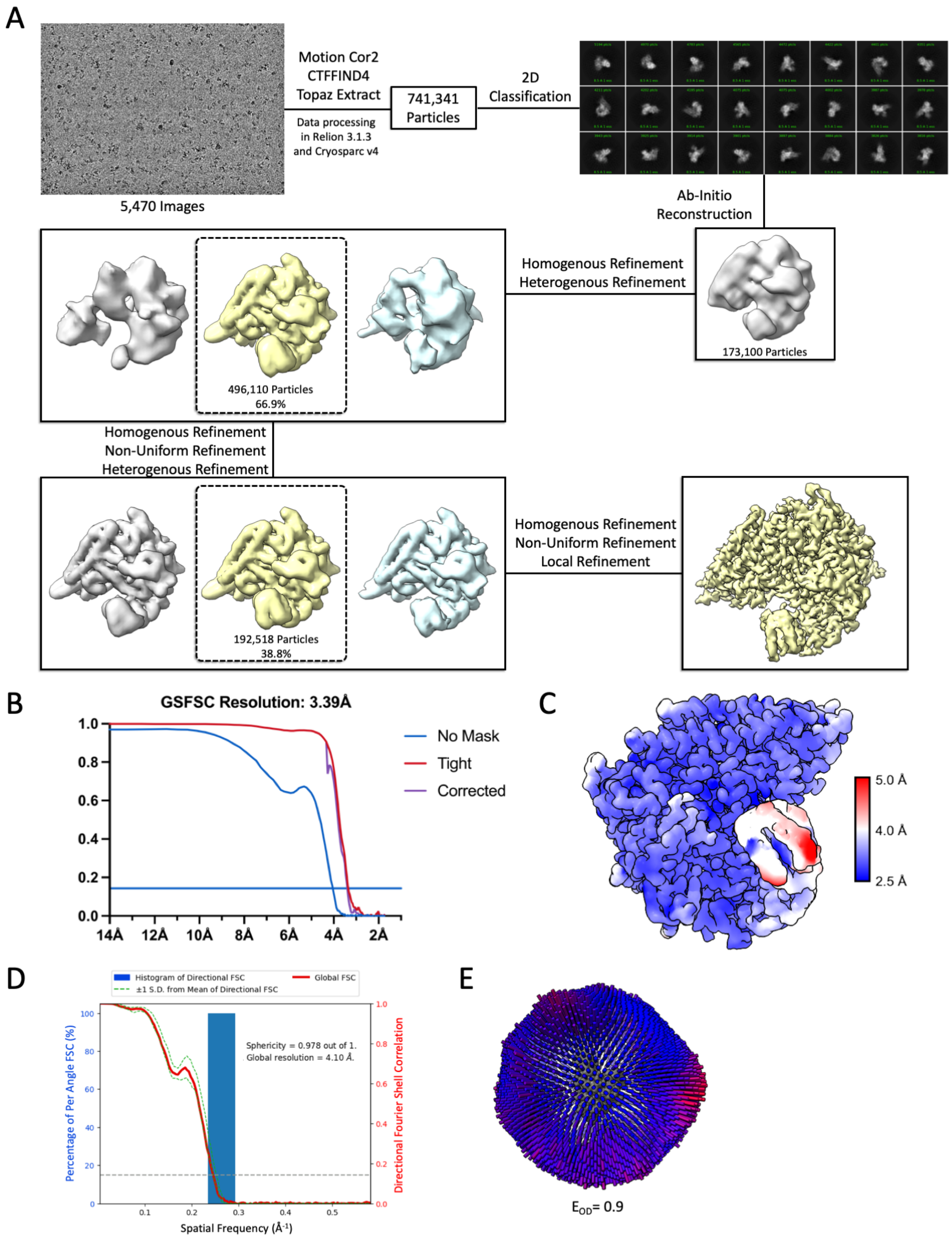

**Figure S7. Cryo-EM workflow and resolution of hNatB/CoA-αS$_{FL}$ structure.** (**A**) Image processing and refinement workflow for hNatB/CoA-αS$_{FL}$ EM map determination. (**B**) Fourier Shell Correlation (FSC) curves of hNatB/CoA-αS$_{FL}$ EM map 3D reconstruction. (**C**) Local resolution of hNatB/CoA-αS$_{FL}$ EM map (**D**) 3DFSC analysis for hNatB/CoA-αS$_{FL}$ EM map determination. (**E**) CryoEF Euler angle distribution.

**Table S1. Cryo-EM data collection, refinement and validation statistics.**

| | |
|---|---|
| **Sample** | hNatB/CoA-α$S_{FL}$ |
| **PDB and EMDB codes** | 8G0L, EMDB-29657 |
| **Data collection and processing** | |
| Magnification | 105,000 |
| Voltage (kV) | 300 |
| Electron exposure (e–/Å$^2$) | 43.9 |
| Defocus range (μm) | -1.0 to -3.0 |
| Pixel size (Å) | 0.43 |
| Symmetry imposed | C1 |
| Initial particle images (no.) | 5,470 |
| Final particle images (no.) | 4,046 |
| Map resolution (Å) | 3.39 |
| FSC threshold | 0.143 |
| Map resolution range (Å) | 3.04-5.20 |
| **Refinement** | |
| Initial model used (PDB code) | 6VP6 |
| Model resolution (Å) | 3.36 |
| FSC threshold | 0.143 |
| Model composition | |
| Non-hydrogen atoms | 8,872 |
| Protein residues | 1,088 |
| Ligands | 1 |
| *B* factors (Å$^2$) | |
| Protein | 176.02 |
| Ligand | 188.24 |
| R.m.s. deviations | |
| Bond lengths (Å) | 0.25 |
| Bond angles (°) | 0.45 |
| Validation | |
| MolProbity score | 2.21 |
| Clashscore | 5 |
| Poor rotamers (%) | 0.1 |
| Ramachandran plot | |
| Favored (%) | 98.51 |
| Allowed (%) | 1.49 |
| Disallowed (%) | 0.00 |

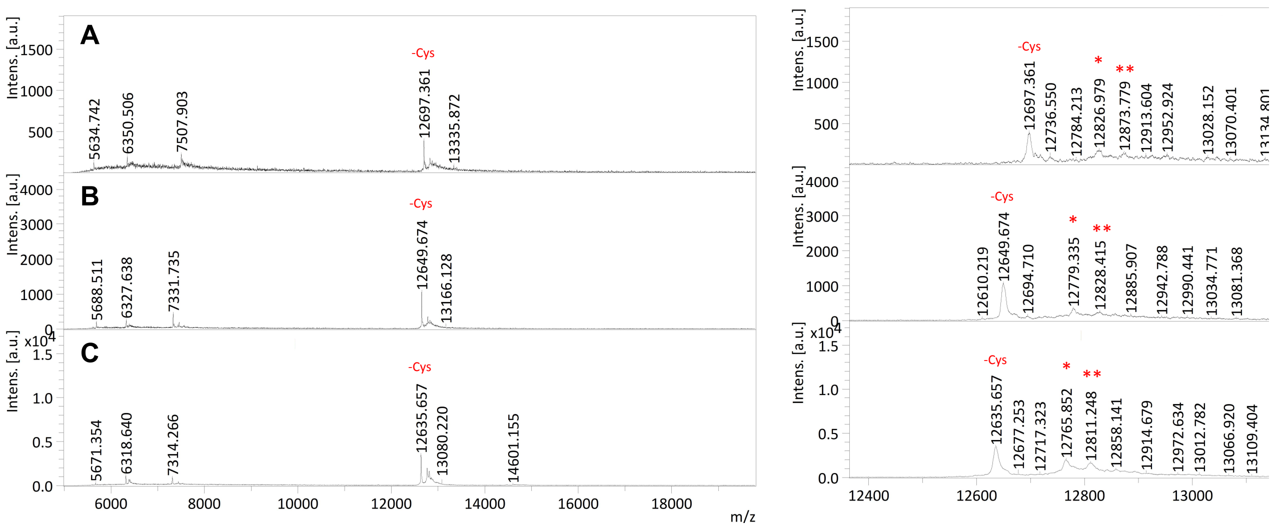

**Figure S8. Recombinant $\alpha S_{18\text{-}140}$-$C_{18}\pi_n$ (S3a, S3b, S3c). (A)** MALDI-MS of recombinant $\alpha S_{18\text{-}140}$-$C_{18}\pi_{72}$. For fragment with free N-terminal cysteine, Calc$[M+H]^+$ = 12801, Obs$[M+H]^+$ = 12697.3. **(B)** MALDI-MS of recombinant $\alpha S_{18\text{-}140}$-$C_{18}\pi_{94}$. For fragment with free N-terminal cysteine, Calc$[M+H]^+$ = 12755, Obs$[M+H]^+$ = 12649.7. **(C)** MALDI-MS of recombinant $\alpha S_{18\text{-}140}$-$C_{18}\pi_{136}$. For fragment with free N-terminal cysteine, Calc$[M+H]^+$ = 12739, Obs$[M+H]^+$ = 12635.7. -Cys = fragment missing cysteine, * = fragment with N-terminal thiazolidine from acetaldehyde ($^{Acet}$Thz), ** = fragment with N-terminal Thz from pyruvate ($^{Pyr}$Thz).

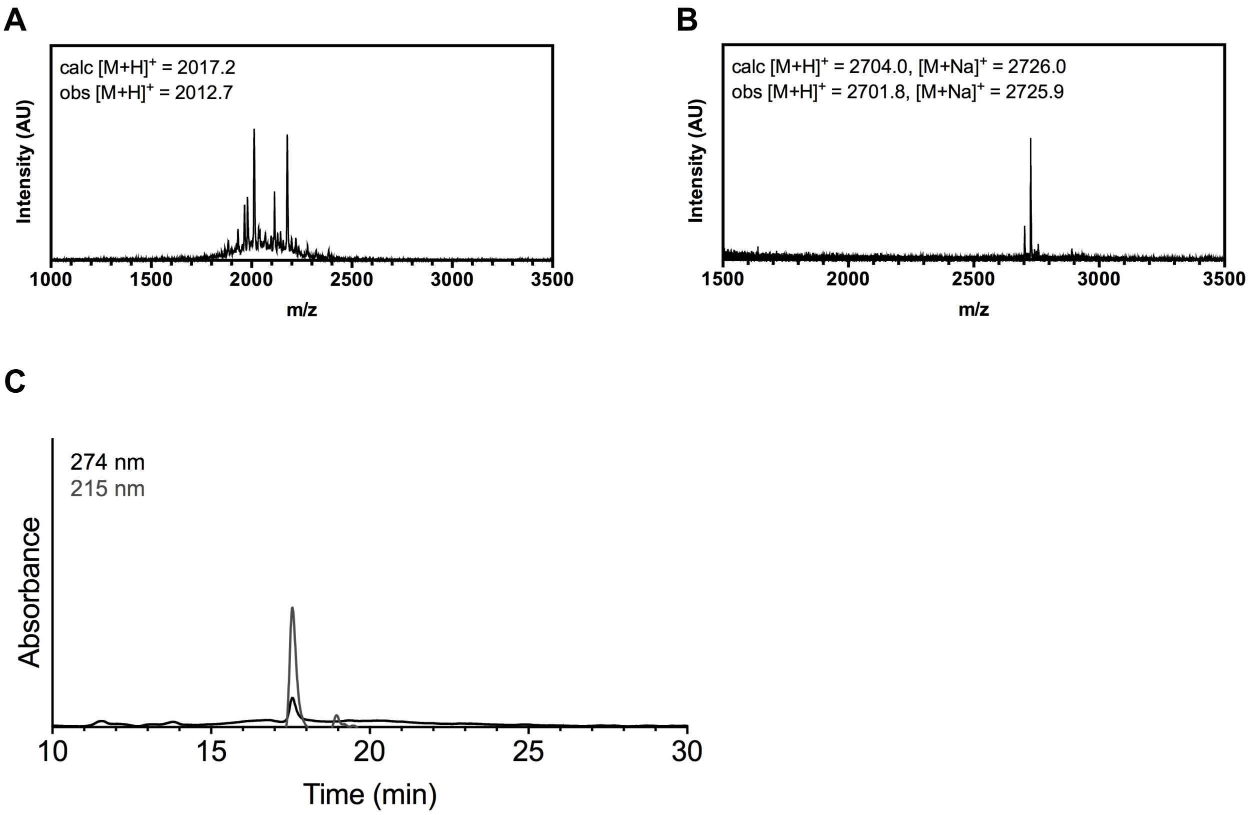

**Figure S9. Synthetic peptide CoA-α$S_{1\text{-}18}$ with C-terminal hydrazide (S2). (A)** MALDI MS of crude Br-Ac-α$S_{1\text{-}18}$, **S1'**. **(B)** MALDI MS of purified CoA-α$S_{1\text{-}18}$ **(C)** Analytical HPLC (10-50% B over 30 min) of purified CoA-α$S_{1\text{-}18}$ (Retention time 17.6 min).

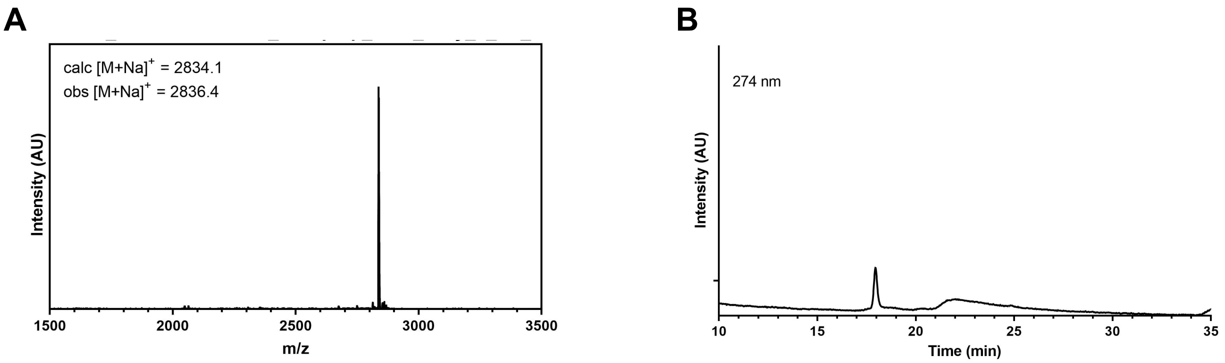

**Figure S10. Synthetic peptide CoA-αS$_{1\text{-}18}$ with C-terminal alkyl thioester (S3). (A)** MALDI MS of purified CoA-αS$_{1\text{-}18}$-S$(CH_2)_2SO_3H$ **(B)** Analytical HPLC (10-50% B over 30 min) of purified CoA-αS$_{1\text{-}18}$-S$(CH_2)_2SO_3H$ (Retention time 18.0 min).

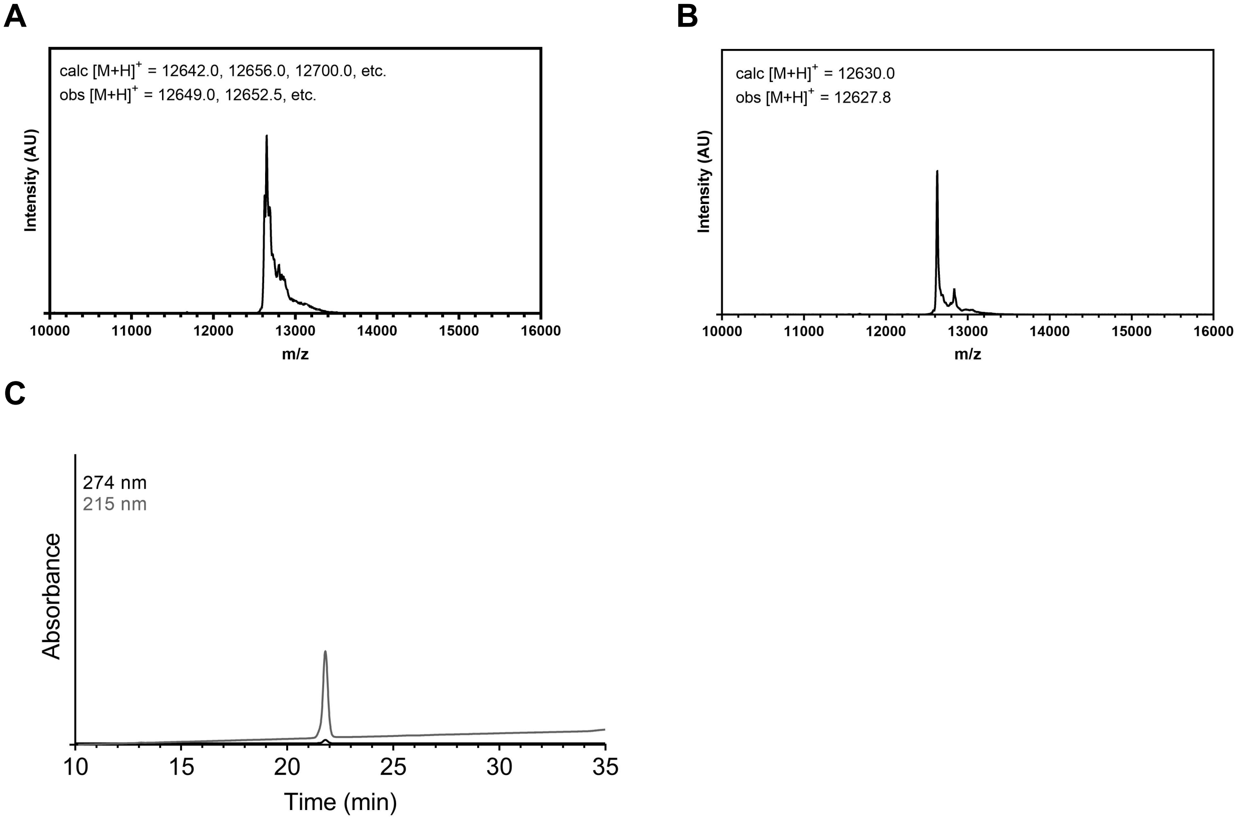

**Figure S11. Recombinant αS$_{19\text{-}140}$-C$_{19}$ (S4). (A)** MALDI MS of Thz-αS$_{19\text{-}140}$-C$_{19}$ starting material. Observed masses correspond to various aldehyde- and ketone- derived thiazolidines. **(B)** MALDI MS of purified product after methoxyamine-mediated deprotection reaction. **(C)** Analytical HPLC (10-60% B over 30 min) of purified αS$_{19\text{-}140}$-C$_{19}$ **S4** (Retention time 21.8 min).

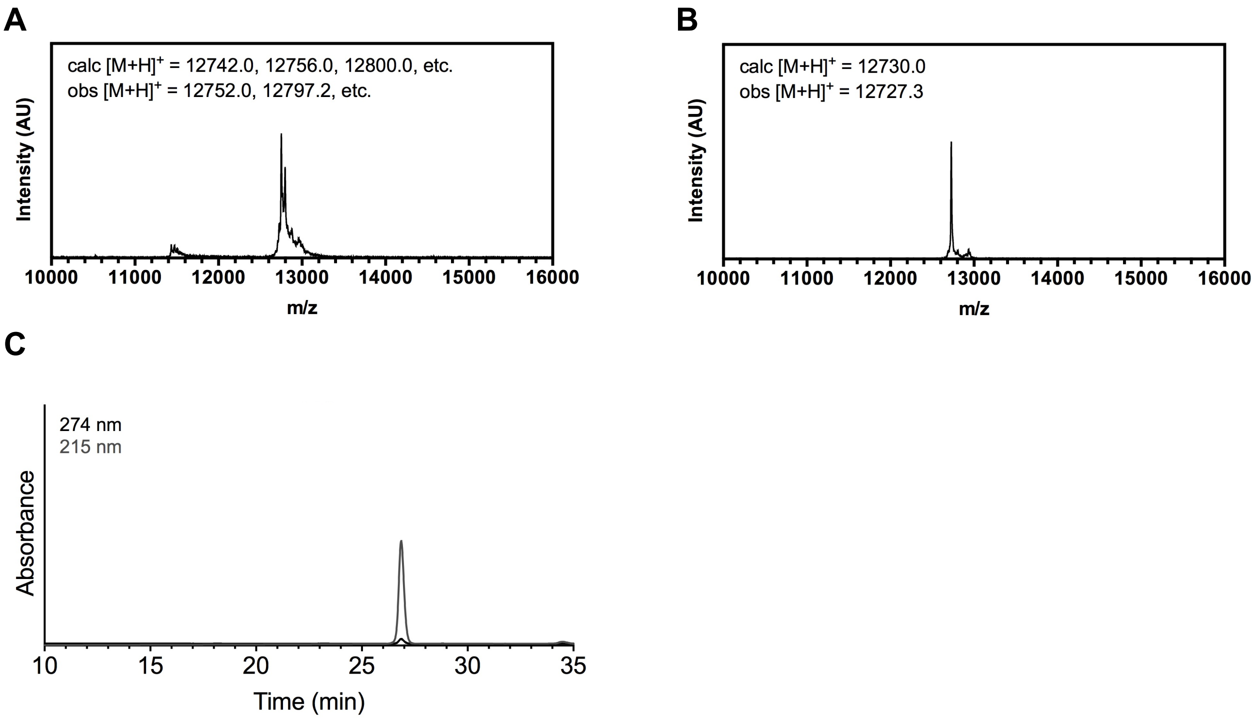

**Figure S12. Recombinant αS$_{19\text{-}140}$-C$_{19}$π$_{72}$ (11a). (A)** MALDI MS of Thz-αS$_{19\text{-}140}$-C$_{19}$π$_{72}$ starting material. Observed masses correspond to various aldehyde- and ketone- derived thiazolidines. **(B)** MALDI MS of purified product after methoxyamine-mediated deprotection reaction. **(C)** Analytical HPLC (10-50% B over 30 min) of purified αS$_{19\text{-}140}$-C$_{19}$π$_{72}$ **11a** (Retention time 26.9 min).

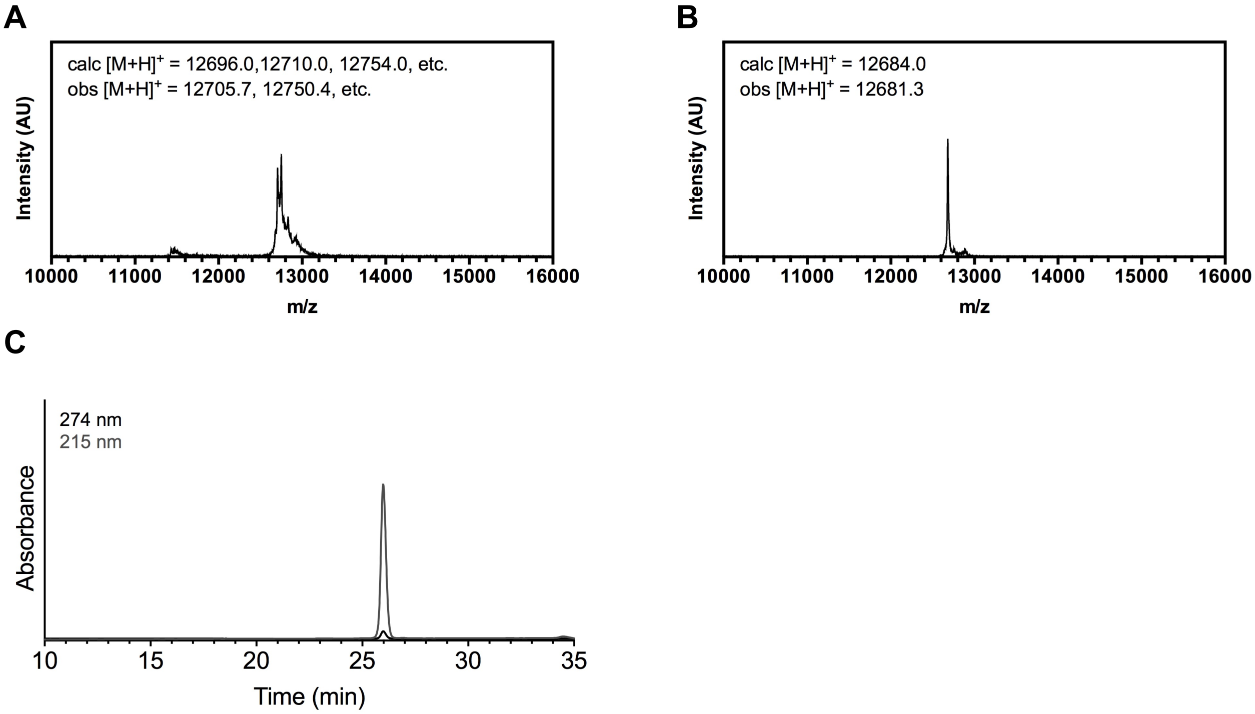

**Figure S13. Recombinant $\alpha S_{19\text{-}140}$-$C_{19}\pi_{94}$ (11b). (A)** MALDI MS of Thz-$\alpha S_{19\text{-}140}$-$C_{19}\pi_{94}$ starting material. Observed masses correspond to various aldehyde- and ketone- derived thiazolidines. **(B)** MALDI MS of purified product after methoxyamine-mediated deprotection reaction. **(C)** Analytical HPLC (10-50% B over 30 min) of purified $\alpha S_{19\text{-}140}$-$C_{19}\pi_{94}$ **11b** (Retention time 26.0 min).

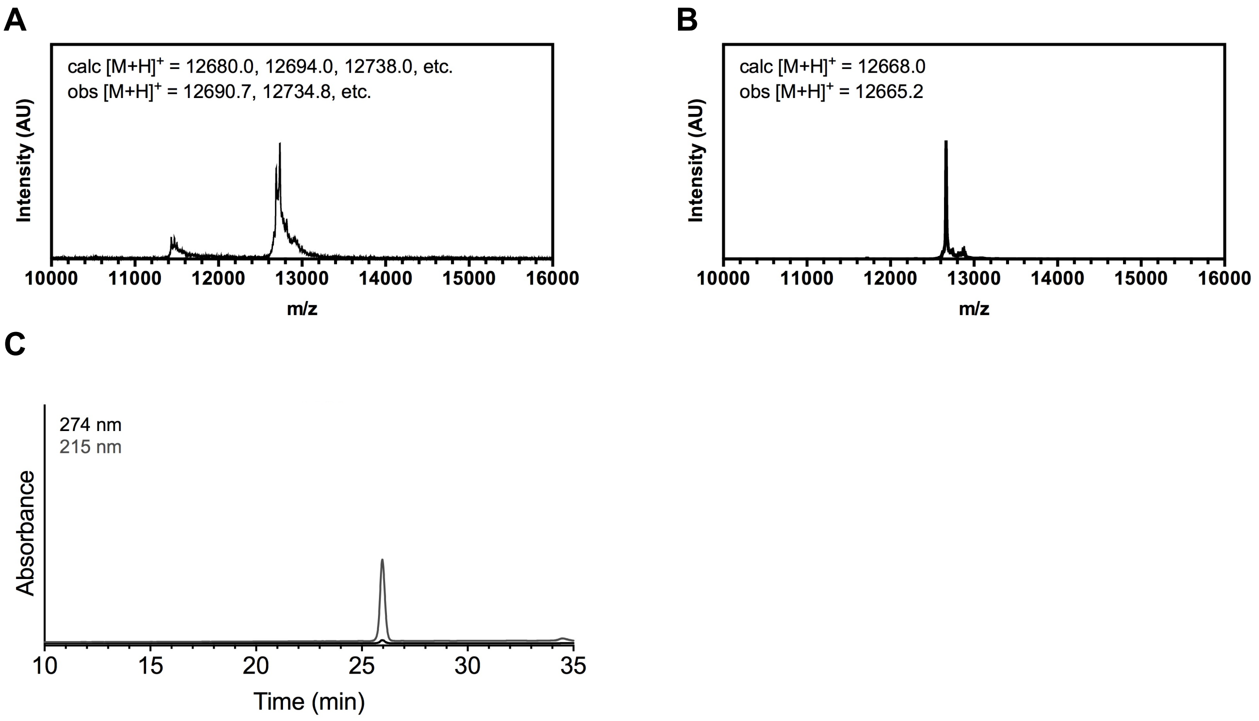

**Figure S14. Recombinant αS$_{19\text{-}140}$-C$_{19}$π$_{136}$ (11c). (A)** MALDI MS of Thz-αS$_{19\text{-}140}$-C$_{19}$π$_{136}$ starting material. Observed masses correspond to various aldehyde- and ketone- derived thiazolidines. **(B)** MALDI MS of purified product after methoxyamine-mediated deprotection reaction. **(C)** Analytical HPLC (10-50% B over 30 min) of purified αS$_{19\text{-}140}$-C$_{19}$π$_{136}$ **11c** (Retention time 26.0 min).

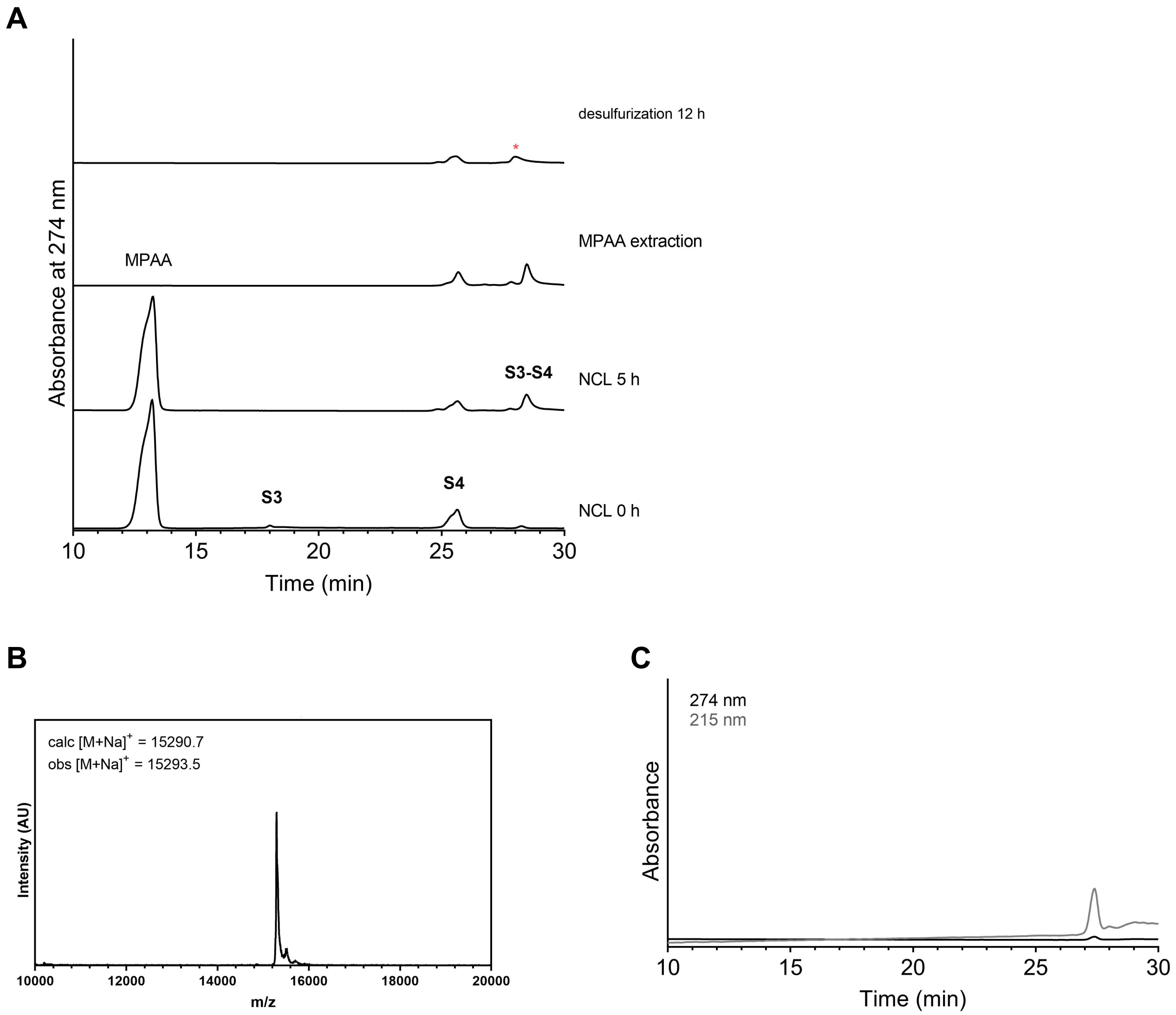

**Figure S15. NCL of CoA-αS$_{1-18}$-MES (S3) and αS$_{19-140}$-C$_{19}$ (S4). (A)** Analytical HPLC (10-50% B over 30 min) of NCL and subsequent desulfurization reaction; CoA-αS$_{1-18}$ (**S3**), αS$_{19-140}$-C$_{19}$π$_{136}$ (**S4**), and product (*) **(B)** MALDI MS of NCL followed by desulfurization product **(C)** Analytical HPLC (10-50% B over 30 min) of purified CoA-αS$_{FL}$ **9** (Retention time 27.4 min).

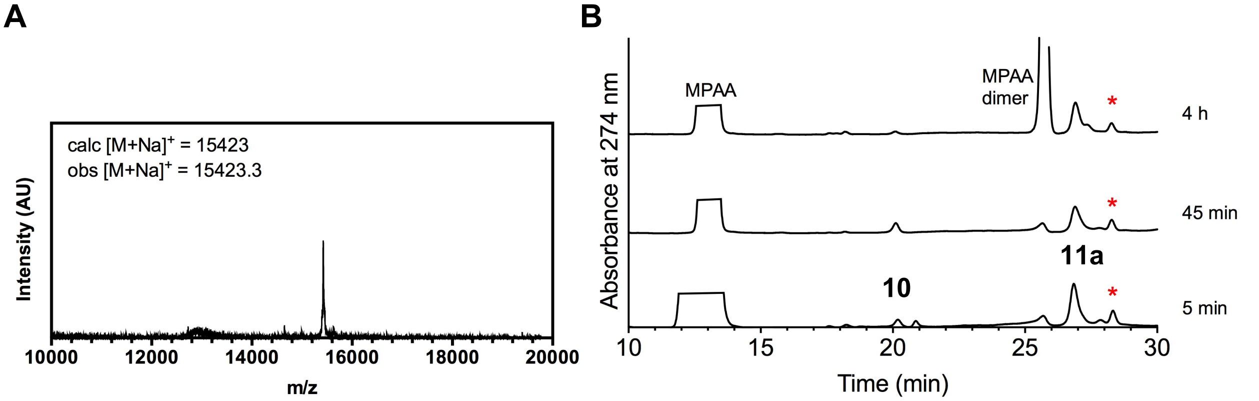

**Figure S16. NCL of CoA-$\alpha S_{1\text{-}18}$ (10) and $\alpha S_{19\text{-}140}$-$C_{19}\pi_{72}$ (11a). (A)** MALDI MS of NCL product **(B)** Analytical HPLC (10-50% B over 30 min) of NCL reaction; CoA-$\alpha S_{1\text{-}18}$ (**10**), $\alpha S_{19\text{-}140}$-$C_{19}\pi_{72}$ (**11a**), and product (*) (Retention time 28.3 min).

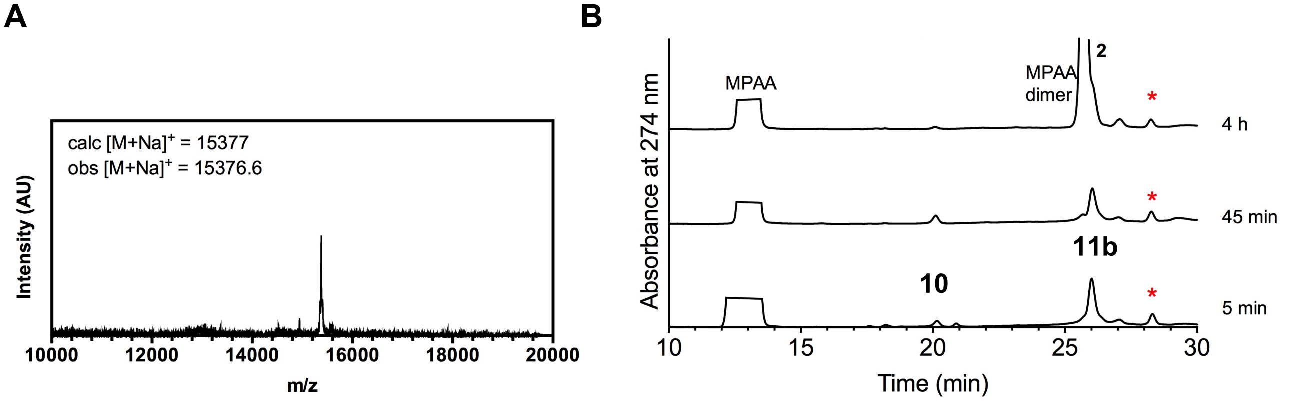

**Figure S17. NCL of CoA-$\alpha S_{1\text{-}18}$ (10) and $\alpha S_{19\text{-}140}$-$C_{19}\pi_{94}$ (11b). (A)** MALDI MS of NCL product **(B)** Analytical HPLC (10-50% B over 30 min) of NCL reaction; CoA-$\alpha S_{1\text{-}18}$ (**10**), $\alpha S_{19\text{-}140}$-$C_{19}\pi_{94}$ (11**b**), and product (*) (Retention time 28.3 min).

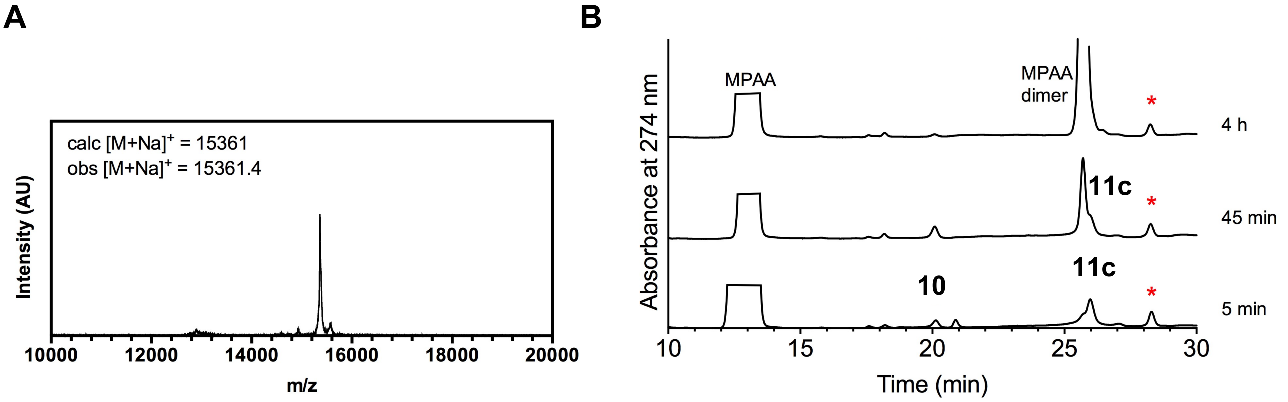

**Figure S18. NCL of CoA-αS$_{1-18}$ (10) and αS$_{19-140}$-C$_{19}$π$_{136}$ (11c). (A)** MALDI MS of NCL product **(B)** Analytical HPLC (10-50% B over 30 min) of NCL reaction; CoA-αS$_{1-18}$ (**a**), αS$_{19-140}$-C$_{19}$π$_{136}$ (**b**), and product (*) (Retention time 28.3 min).

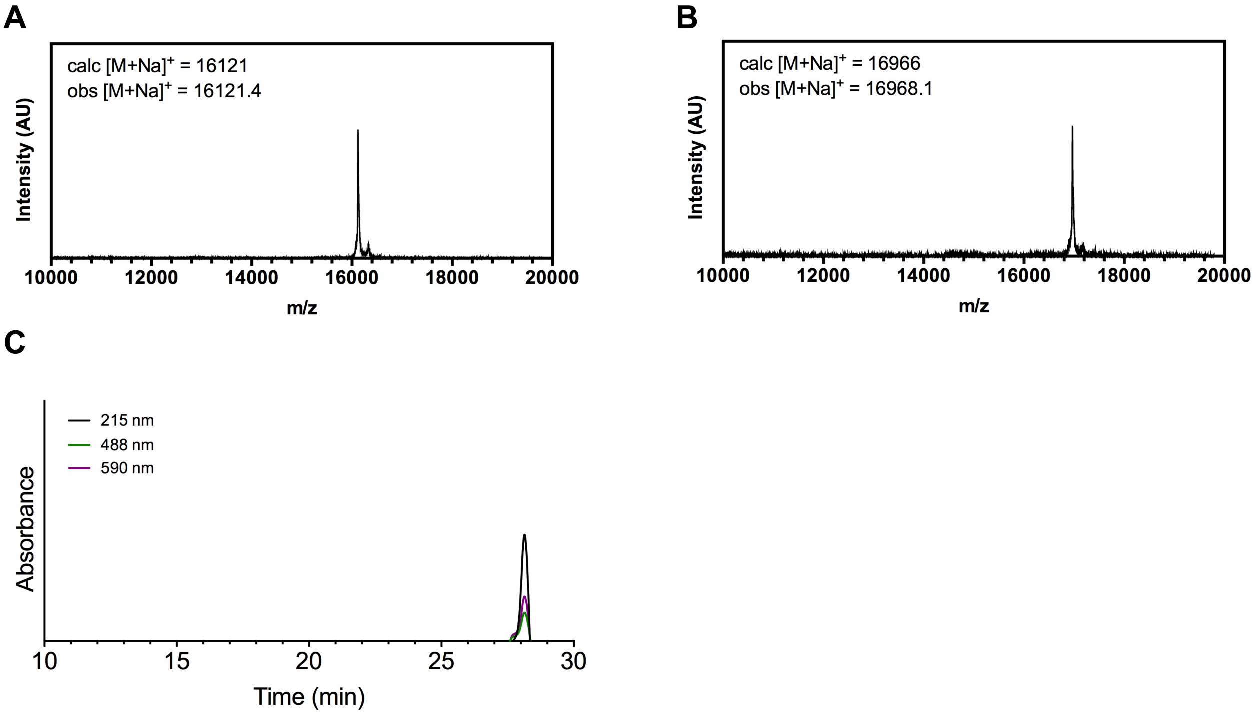

**Figure S19. Fluorescent labeling of CoA-αS-C$_{19}$π$_{72}$ (12a). (A)** MALDI MS of AF488-maleimide labeled CoA-αS-C$^{488}_{19}$π$_{72}$ **(B)** MALDI MS of AF594-N$_3$ labeled CoA-αS-C$^{488}_{19}$π$^{594}_{72}$ **(C)** Analytical HPLC (10-50% B over 30 min) of purified CoA-αS-C$^{488}_{19}$π$^{594}_{72}$ (Retention time 28.0 min).

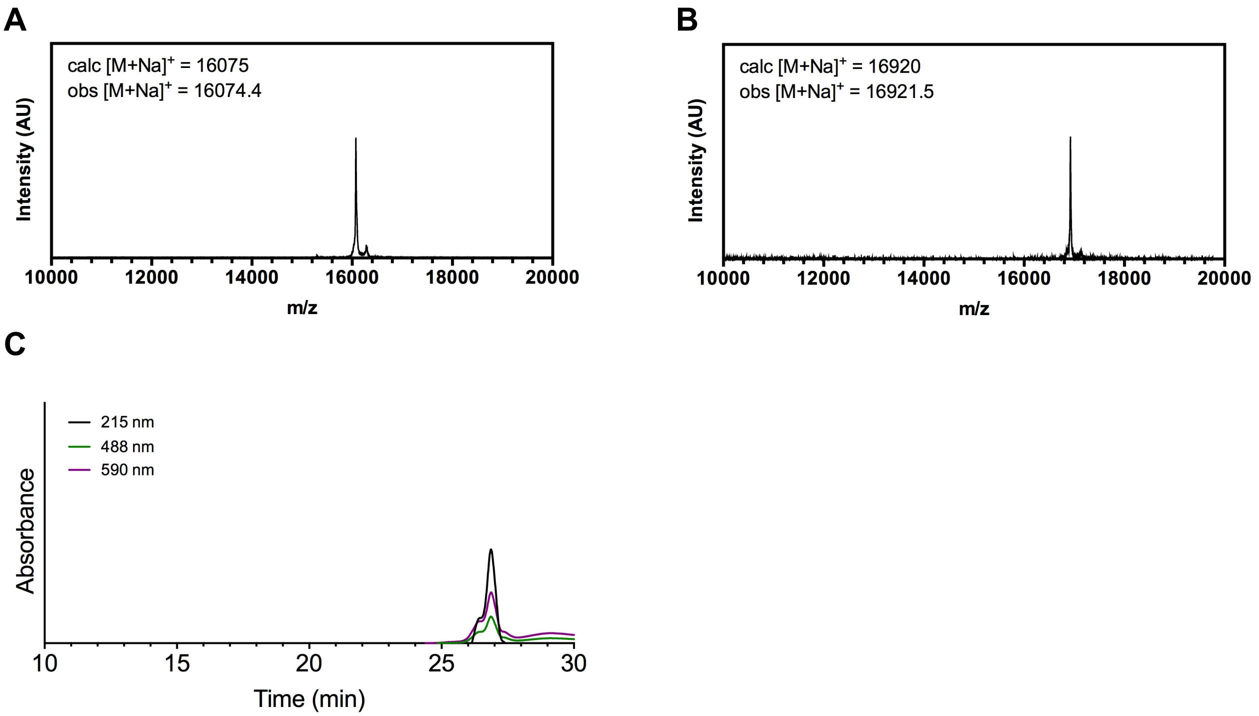

**Figure S20. Fluorescent labeling of CoA-αS-C$_{19}$π$_{94}$ (12b). (A)** MALDI MS of AF488-maleimide labeled CoA-αS-C$^{488}_{19}$π$_{94}$ **(B)** MALDI MS of AF594-$N_3$ labeled CoA-αS-C$^{488}_{19}$π$^{594}_{94}$ **(C)** Analytical HPLC (10-50% B over 30 min) of purified CoA-αS-C$^{488}_{19}$π$^{594}_{94}$ (Retention time 26.9 min).

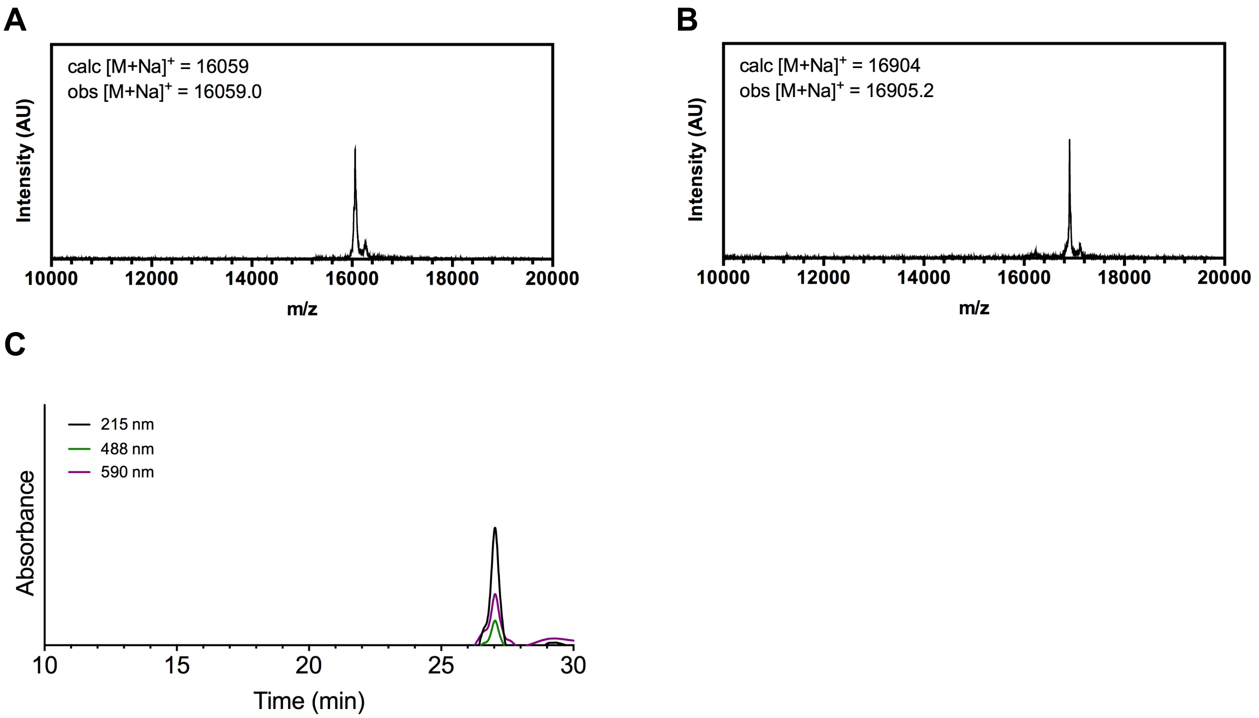

**Figure S21. Fluorescent labeling of CoA-αS-$C_{19}\pi_{136}$ (12c). (A)** MALDI MS of AF488-maleimide labeled CoA-αS-$C^{488}{}_{19}\pi_{136}$ **(B)** MALDI MS of AF594-$N_3$ labeled CoA-αS-$C^{488}{}_{19}\pi^{594}{}_{136}$ **(C)** Analytical HPLC (10-50% B over 30 min) of purified CoA-αS-$C^{488}{}_{19}\pi^{594}{}_{136}$ (Retention time 27.1 min).

**Table S2. FCS data.**

| Sample | Diffusion Time (ms) |
|---|---|
| CoA-αS-$C^{488}{}_{19}\pi$ free in solution | 0.504 ± 0.005 |
| CoA-αS-$C^{488}{}_{19}\pi$ with 1 equiv NatB | 0.543 ± 0.002 |
| CoA-αS-$C^{488}{}_{19}\pi$ with 10 equiv NatB | 0.704 ± 0.002 |
| CoA-αS-$C^{488}{}_{19}\pi$ with 40 equiv NatB | 0.854 ± 0.007 |
| CoA-αS-$C^{488}{}_{19}\pi$ with 100 equiv NatB | 0.858 ± 0.004 |

The diffusion time of each sample was determined by fitting the autocorrelation curves to a single diffusing component. Results shown as mean with standard deviation (n=3)

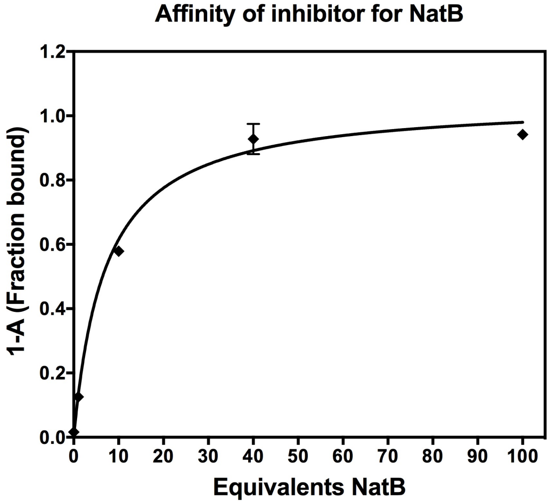

**Figure S22. Full binding of CoA-αS-$C^{488}{}_{19}\pi_n$ (13a, 13b, 13c) to NatB by FCS.** The fraction of inhibitor bound to NatB at each molar equivalents of the enzyme was determined by fitting the autocorrelation curves to a two-component equation (see Methods).

**Table S3. smFRET data.**

| | $ET_{eff}$ | | | Mean $ET_{eff}$ | Stdev $ET_{eff}$ |
|---|---|---|---|---|---|
| CoA-αS-$C^{488}{}_{19}\pi^{594}{}_{72}$ (**13a**) | 0.65 | 0.65 | 0.65 | 0.65 | 0.00 |
| NatB-CoA-αS-$C^{488}{}_{19}\pi^{594}{}_{72}$ (**13a**) | 0.65 | 0.65 | 0.64 | 0.65 | 0.01 |
| CoA-αS-$C^{488}{}_{19}\pi^{594}{}_{94}$ (**13b**) | 0.48 | 0.48 | 0.48 | 0.48 | 0.00 |
| NatB-CoA-αS-$C^{488}{}_{19}\pi^{594}{}_{94}$ (**13b**) | 0.42 | 0.43 | 0.44 | 0.43 | 0.01 |
| CoA-αS-$C^{488}{}_{19}\pi^{594}{}_{136}$ (**13c**) | 0.30 | 0.30 | 0.30 | 0.30 | 0.00 |
| NatB-CoA-αS-$C^{488}{}_{19}\pi^{594}{}_{136}$ (**13c**) | 0.24 | 0.25 | 0.25 | 0.25 | 0.01 |

| | Width | | | Mean width | Stdev width |
|---|---|---|---|---|---|
| CoA-αS-$C^{488}{}_{19}\pi^{594}{}_{72}$ (**13a**) | 0.15 | 0.15 | 0.16 | 0.15 | 0.01 |
| NatB-CoA-αS-$C^{488}{}_{19}\pi^{594}{}_{72}$ (**13a**) | 0.17 | 0.17 | 0.17 | 0.17 | 0.00 |
| CoA-αS-$C^{488}{}_{19}\pi^{594}{}_{94}$ (**13b**) | 0.16 | 0.17 | 0.16 | 0.16 | 0.01 |
| NatB-CoA-αS-$C^{488}{}_{19}\pi^{594}{}_{94}$ (**13b**) | 0.23 | 0.24 | 0.25 | 0.24 | 0.01 |
| CoA-αS-$C^{488}{}_{19}\pi^{594}{}_{136}$ (**13c**) | 0.20 | 0.21 | 0.21 | 0.21 | 0.01 |
| NatB-CoA-αS-$C^{488}{}_{19}\pi^{594}{}_{136}$ (**13c**) | 0.24 | 0.25 | 0.22 | 0.24 | 0.01 |

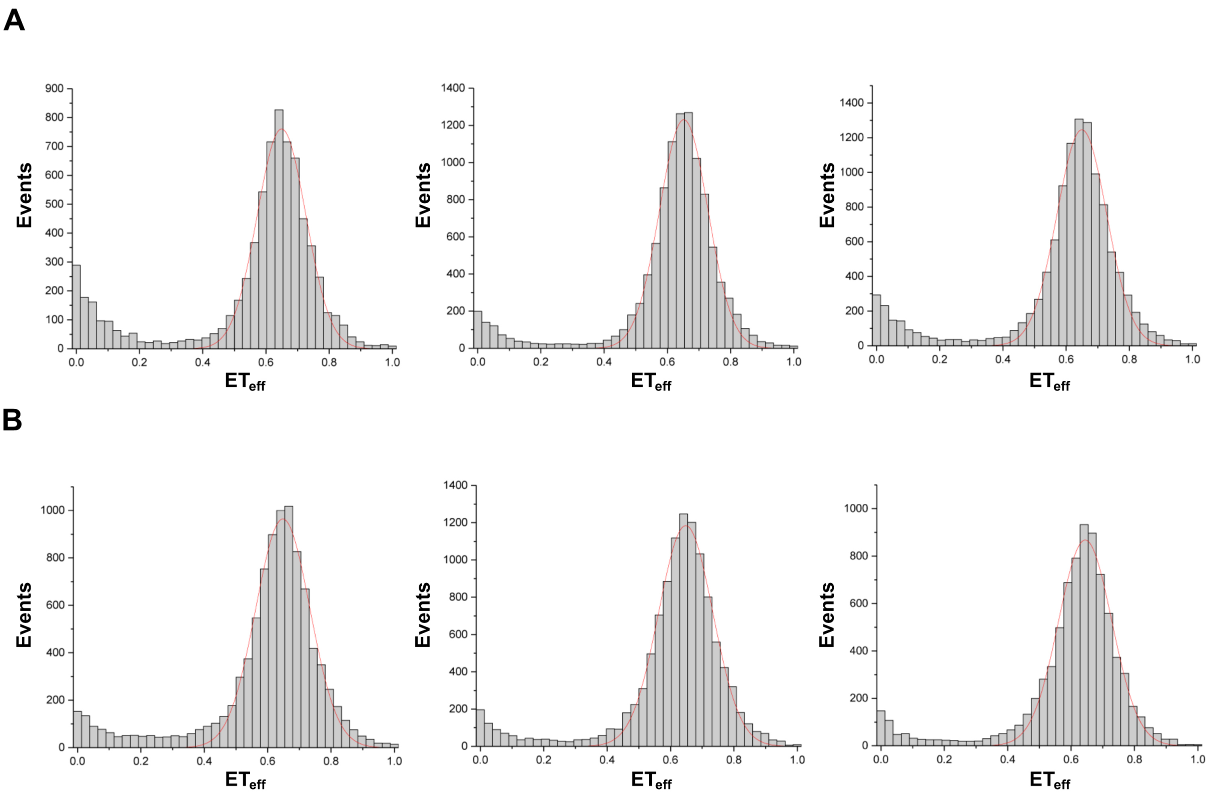

**Figure S23. Conformation of CoA-αS-$C^{488}{}_{19}\pi^{594}{}_{72}$ (13a) and complex with NatB by smFRET.** smFRET histograms comparing **(A)** CoA-αS $C^{488}{}_{19}\pi^{594}{}_{72}$ inhibitor alone and **(B)** in complex with hNatB. All data were fit to Gaussian distributions.

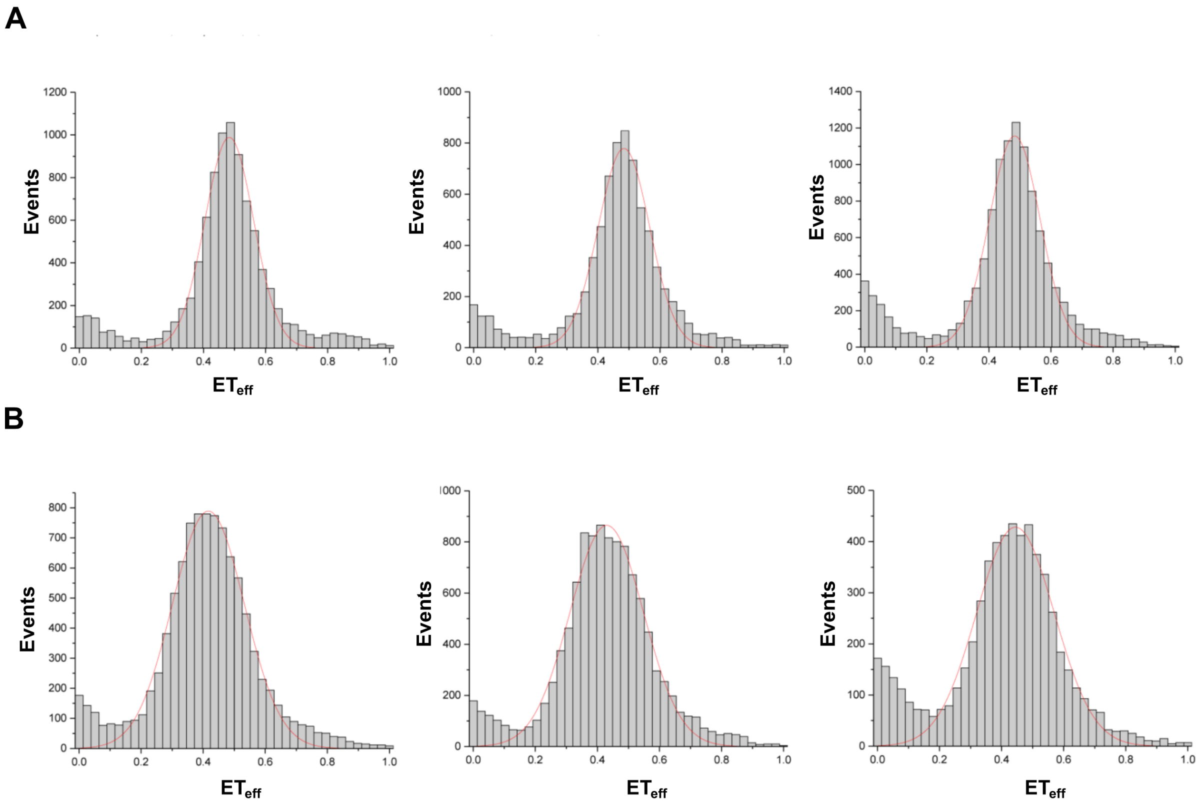

**Figure S24. Conformation of CoA-αS-$C^{488}{}_{19}\pi^{594}{}_{94}$ (13b) and complex with NatB by smFRET.** smFRET histograms comparing **(A)** CoA-αS $C^{488}{}_{19}\pi^{594}{}_{94}$ inhibitor alone and **(B)** in complex with hNatB. All data were fit to Gaussian distributions.

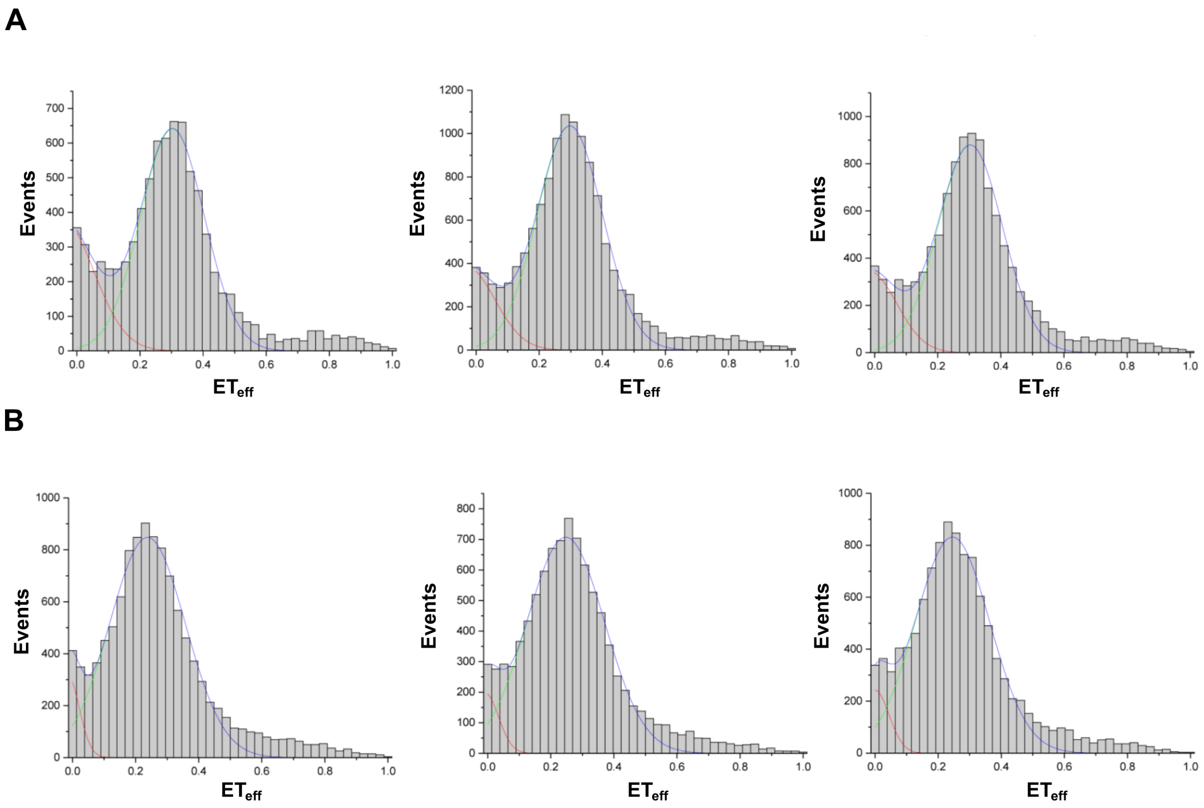

**Figure S25. Conformation of CoA-αS-$C^{488}{}_{19}\pi^{594}{}_{136}$ (13c) and complex with NatB by smFRET.** smFRET histograms comparing **(A)** CoA-αS $C^{488}{}_{19}\pi^{594}{}_{136}$ inhibitor alone and **(B)** in complex with hNatB. All data were fit to Gaussian distributions.

**Table S4. Fluorescence anisotropy data.**

| | Anisotropy | | | Mean | Stdev |
|---|---|---|---|---|---|
| CoA-αS-$C^{488}{}_{19}\pi^{594}{}_{72}$ (**13a**) | 0.12 | 0.12 | 0.12 | 0.12 | 0.00 |
| NatB-CoA-αS-$C^{488}{}_{19}\pi^{594}{}_{72}$ (**13a**) | 0.17 | 0.19 | 0.18 | 0.18 | 0.01 |
| CoA-αS-$C^{488}{}_{19}\pi^{594}{}_{94}$ (**13b**) | 0.11 | 0.11 | 0.10 | 0.11 | 0.01 |
| NatB-CoA-αS-$C^{488}{}_{19}\pi^{594}{}_{94}$ (**13b**) | 0.14 | 0.14 | 0.12 | 0.13 | 0.01 |
| CoA-αS-$C^{488}{}_{19}\pi^{594}{}_{136}$ (**13c**) | 0.10 | 0.12 | 0.10 | 0.11 | 0.01 |
| NatB-CoA-αS-$C^{488}{}_{19}\pi^{594}{}_{136}$ (**13c**) | 0.14 | 0.13 | 0.13 | 0.13 | 0.01 |

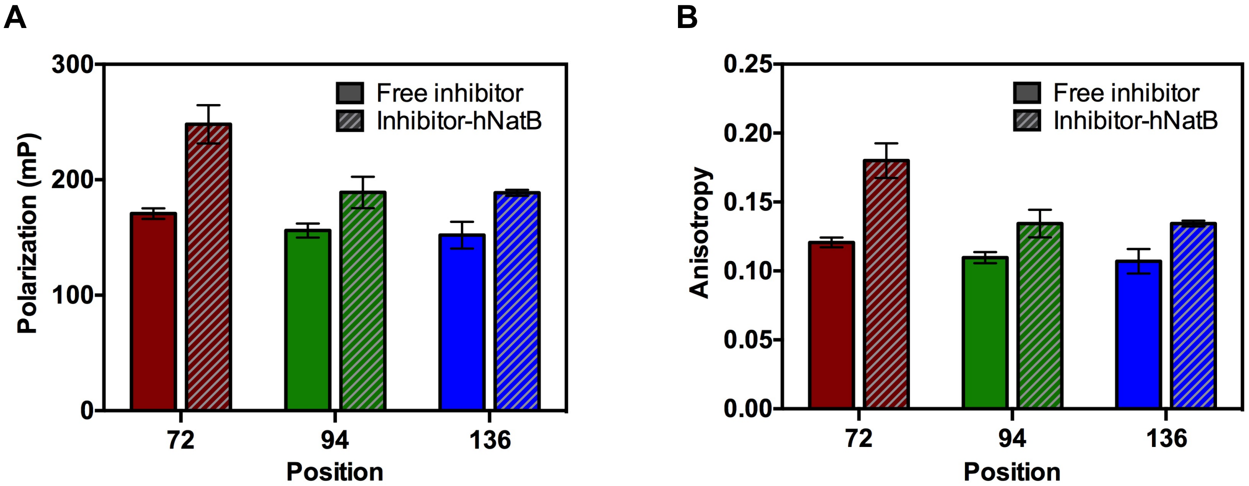

**Figure S26. Fluorescence polarization/anisotropy of CoA-αS-$C^{488}{}_{19}\pi_n$ (13a, 13b, 13c) ± NatB.** **(A)** Fluorescence polarization and **(B)** anisotropy for CoA-αS labeled with AF594 at positions 72, 94, or 136, free in solution or in complex with hNatB.

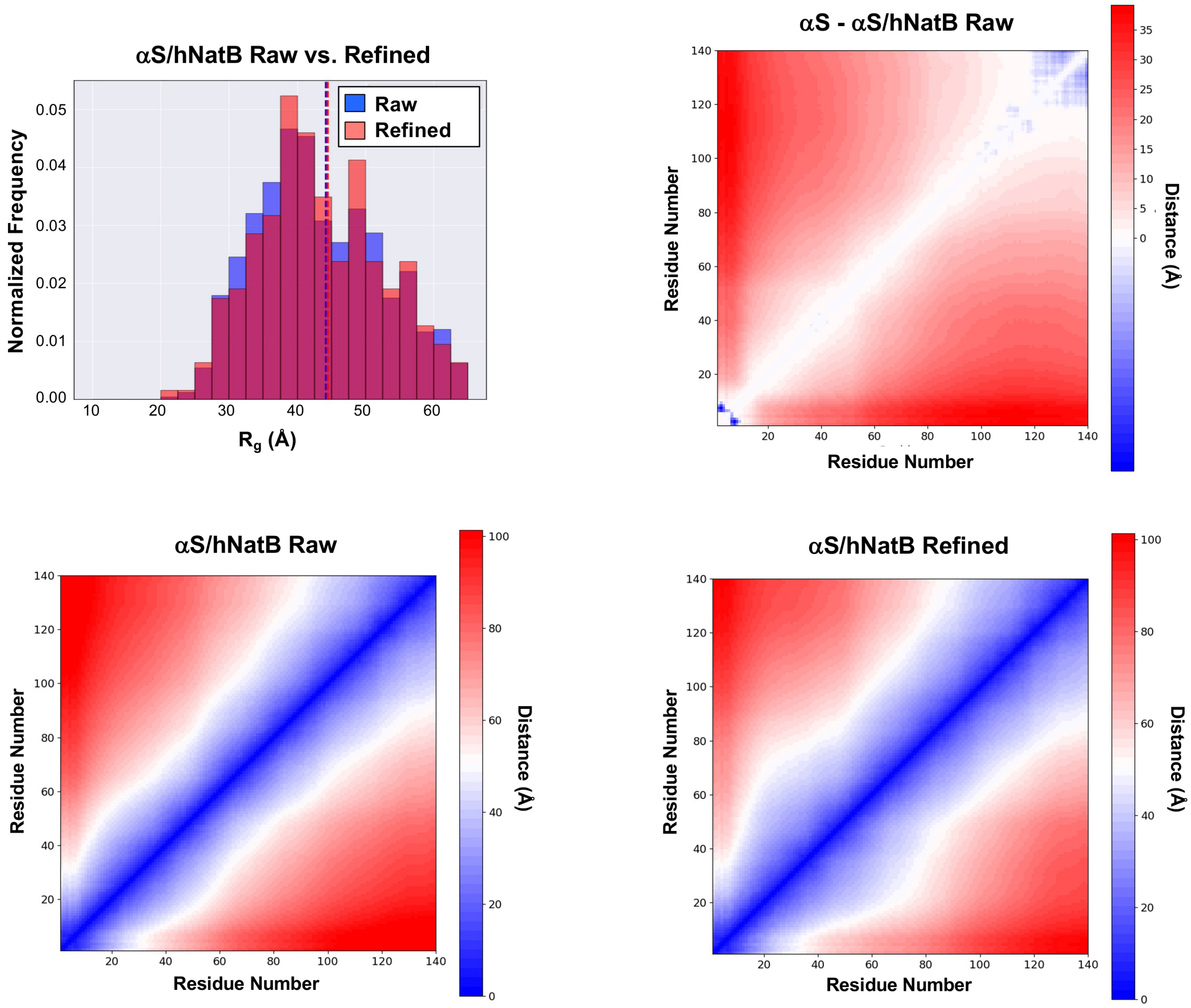

**Figure S27. Analysis of simulated αS/hNatB structures.** Top Left: The radius of gyration (Rg) of the 1000 lowest energy structures in the raw simulated αS/hNatB ensemble (blue) or the 262 structures in the refined αS/hNatB ensemble (red). Dashed lines indicating the ensemble averages are colored accordingly. Top Right: A difference heat map showing the changes in the average inter-residue distances between the αS and raw αS/hNatB ensembles. Bottom: Comparison of contact maps showing inter-residue distances for αS in the ensembles for the raw and refined simulations.

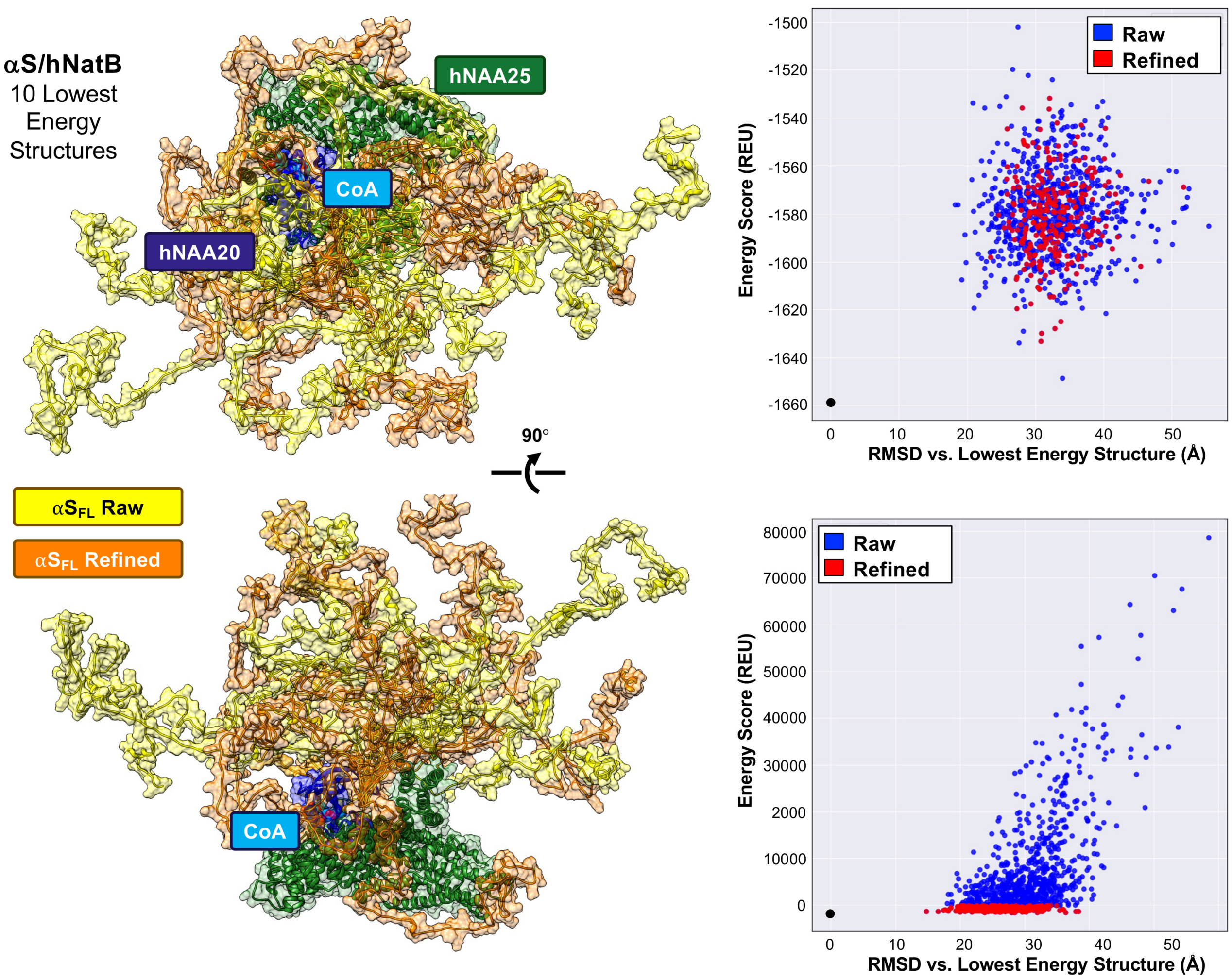

**Figure S28. Representative structures and energies from raw and refined αS/hNatB ensembles.** Left: The CoA-$\alpha S_{FL}$/hNatB complex was simulated in PyRosetta, using the cryo-EM structure as a starting point. The resulting ensemble was then refined using smFRET-derived distance constraints. The 10 lowest energy structures of the αS/hNatB complex from the raw and refined ensembles are overlayed. They are shown in the same two orientations as in main text Figures 2 and 5, with transparent space-filling models superimposed. Right: The energy distributions of the raw (blue) and refined (yellow) structural ensembles are shown with structures organized by root mean squared deviation (RMSD) relative to the lowest energy structure (black). The distributions are shown scored based on the original score function (Top) or the modified score function incorporating distance constraints from the smFRET measurements.

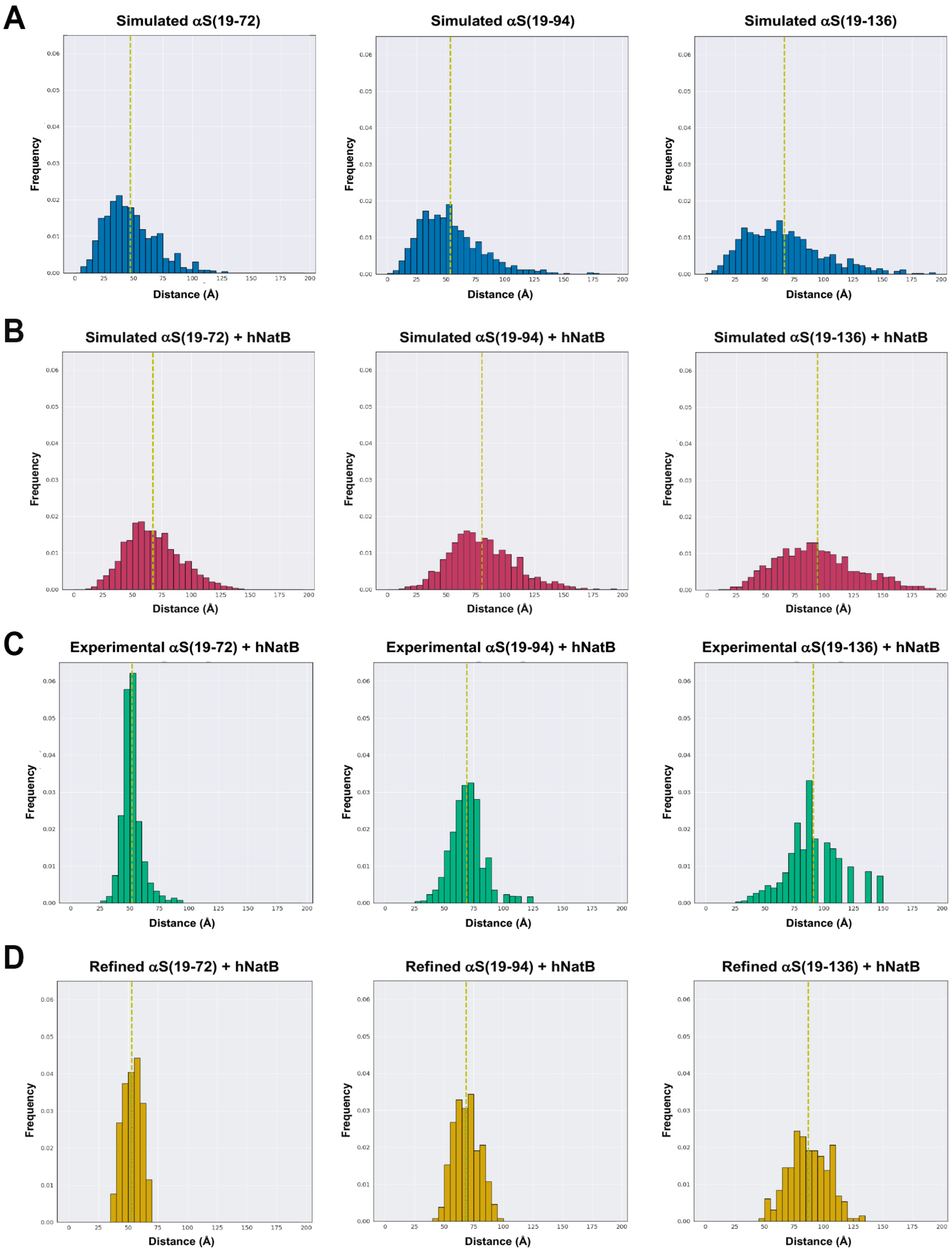

**Figure S29. Simulated and experimental distance distributions for residues in smFRET.** A: Cα-Cα distances of the 1000 lowest energy structures of free αS simulated in Rosetta using FFT as previously described in Ferrie *et al*.[18] B: Cα-Cα distances of the 1000 lowest energy structures of αS in complex with hNatB simulated in Rosetta using FFT with hNatB and αS residues 1-5 held fixed. C: Inter-residue distances derived from smFRET measurements using a Gaussin chain model. αS(19-n) refers to measurements made with CoA-αS-$C^{488}{}_{19}\pi^{594}{}_{n}$. D: Cα-Cα distances of the 262 structures of αS in complex with hNatB after refinement using the smFRET distributions as constraints.

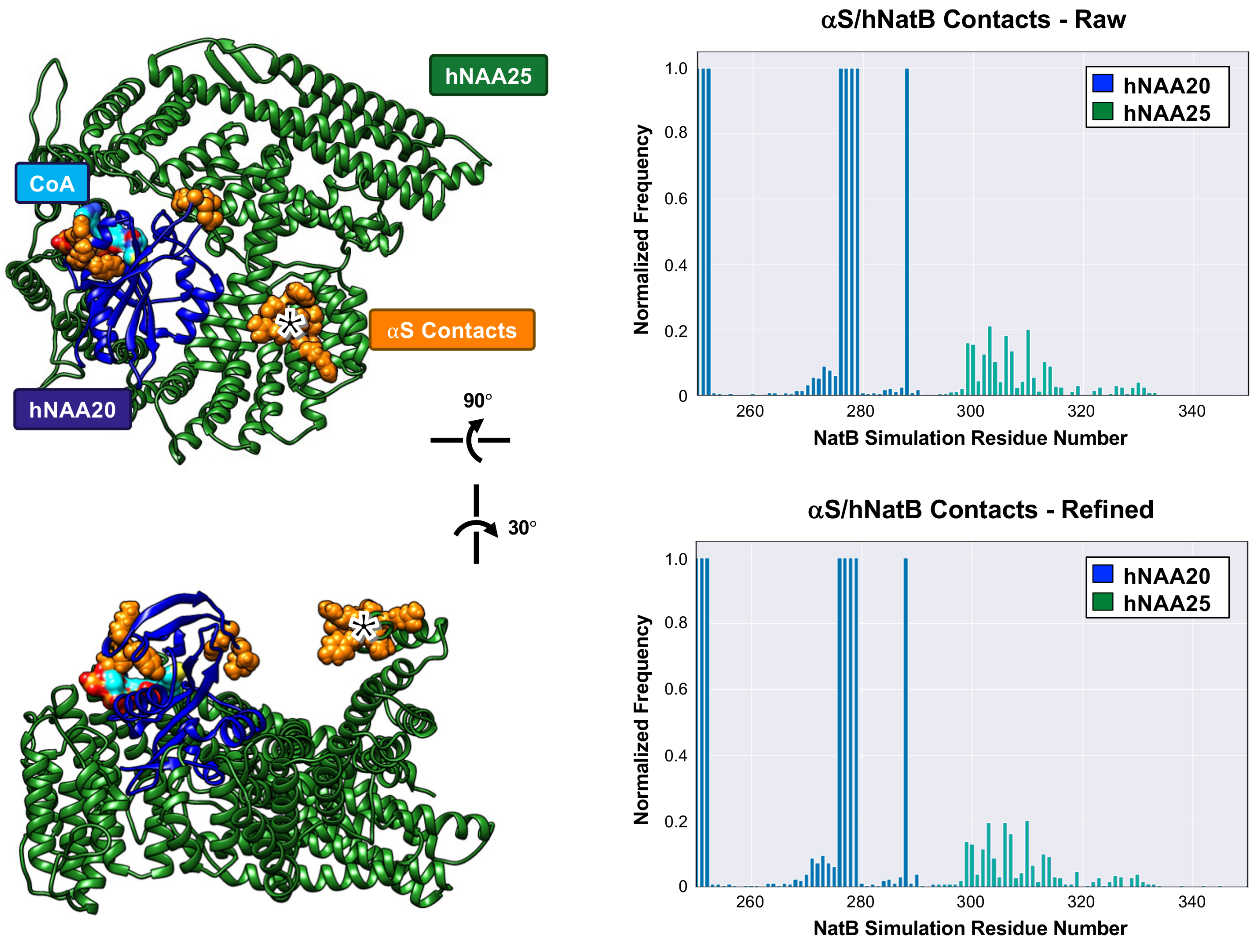

**Figure S30. αS/hNatB contact propensities.** Left: hNatB structure shown in approximately the same two orientations as in main text Figures 2 and 5, with residues that most frequently contact αS indicated as orange spheres. Trivial contacts with the first 5 residues of αS are not highlighted. * indicates hNAA25 residues 13-20. Right: The distributions for structures from the raw (Top) and refined (Bottom) ensembles, determined using a 10 Å cutoff in a minimum of 5% of structures in the ensemble. NatB simulation numbers do not match the sequence of hNAA25 since hNAA20 and hNAA25 were pseudo-concatenated to enable simulation of the αS ensemble.
